# Transformation-tolerant object recognition in tree shrews despite lacking a fovea

**DOI:** 10.64898/2026.04.10.717715

**Authors:** Emily E Meyer, Wei Song Ong, Chenjie Song, Nicolas P Cottaris, Lingqi Zhang, Jared Collina, David H Brainard, Michael J Arcaro

## Abstract

Object recognition depends on the ability to extract stable representations across changes in how they are viewed, yet it remains unclear how this capacity depends on visual acuity and cortical hierarchy. We combined behavioral testing and computational modeling to determine whether tree shrews, close relatives of primates with lower spatial acuity, can perform transformation-tolerant object recognition. Front-end modeling incorporating species-specific optics and photoreceptor sampling showed that, when scaled for acuity, tree shrew retinal filtering preserves the similarity structure of natural image categories relevant for object recognition. Behaviorally, tree shrews reliably discriminated complex objects across variations in position, scale, and viewpoint, including when embedded within natural scenes, and generalized to novel exemplars. Their recognition behavior was best explained by visual features emphasizing differences in global shape and size between objects and by representations from intermediate and deep layers of hierarchical neural network models. These results demonstrate that visual processing supporting object-level generalization can arise within visual systems lacking high-acuity front-end optics and establish the tree shrew as a key model for understanding the computational and evolutionary origins of high-level vision.

## Introduction

A defining feature of human vision is the ability to recognize objects rapidly and accurately despite substantial changes in how they are viewed. An object can shift in position, grow or shrink on the retina as viewing distance changes, rotate to reveal different viewpoints, or appear against varied backgrounds, yet we can perceive it as having the same identity. This capacity, often termed transformation-tolerant or invariant object recognition (Riesenhuber and Poggio, 1999; DiCarlo et al., 2012), is fundamental to how we interact with the visual world. Achieving it depends on both the information captured by the retina and how subsequent stages of the visual system transform that information.

Humans actively exploit a foveated visual system, directing gaze to place objects of interest onto a region of dense photoreceptor sampling that provides high-acuity input to downstream visual areas. Processing of this input is organized hierarchically, with information passing through a series of stages that progressively combine simpler features into more complex representations that are selective for object identity but tolerate variations in appearance (Rust and DiCarlo, 2010; DiCarlo et al., 2012). Models inspired by this cortical architecture, from classical feedforward theoretical frameworks to modern convolutional neural networks, capture this progression of increasing invariance but are typically optimized and evaluated using high-resolution visual input containing fine spatial detail (Riesenhuber and Poggio, 1999; Krizhevsky et al., 2012; Kubilius et al., 2018; Zhuang et al., 2021). This correspondence between high-acuity input and hierarchical computation mirrors the foveal vision of primates and raises a central question: to what extent does transformation-tolerant recognition depend on foveal precision and dense retinal sampling, and to what extent can it arise from hierarchical computation operating on coarser visual input?

Evidence from comparative studies suggests that both optical acuity and cortical hierarchy constrain object recognition abilities. Much of what is known about invariant object recognition comes from studies in humans and other primates, which share similar accuracy and category discriminability patterns (Majaj et al., 2015; Rajalingham et al., 2015; Luongo et al., 2023). Furthermore, humans and other primates have been shown to use only a narrow range of spatial frequencies for fine face, object, and category identification (Näsänen, 1999; Subramanian et al., 2023; Farhang et al., 2025), indicating that object processing may be limited with coarser spatial content. Rodents, by contrast, can perform basic object discrimination, but show more limited tolerance to transformations and reduced performance in complex scenes, consistent both with their low-acuity eyes and comparatively shallow visual hierarchies (Zoccolan et al., 2009; Djurdjevic et al., 2018; Vinken and Beeck, 2021; Schnell et al., 2023a, 2023b). These species differences have been taken to suggest that both high-resolution input and deep hierarchical processing are required for complex object recognition.

Yet empirical findings from degraded visual conditions complicate the view that fine spatial detail is necessary for complex object recognition. Blur is pervasive in everyday human vision due to defocus, peripheral viewing, and the slow dynamics of lens accommodation (Burge and Geisler, 2011), yet humans remain adept at recognizing objects and faces even under substantial blur (Bar, 2004; Goffaux and Rossion, 2006; Kwon and Legge, 2012). Recognition likewise remains robust when stimuli are presented in the lower-acuity periphery, where high spatial frequencies are attenuated, indicating that coarse spatial information can be sufficient for successful discrimination (Thorpe et al., 2001; Larson and Loschky, 2009). Consistent with these behavioral observations, neurons in primate visual cortex explicitly encode blur while maintaining shape-selective responses across changes in image quality (Oleskiw et al., 2018). Computational studies further show that including blurred images during deep neural network training improves robustness and alignment with biological vision (Jang and Tong, 2024). These findings challenge the assumption that high-acuity input is strictly required for complex object recognition and highlight that comparisons limited to primates and rodents may confound the roles of retinal resolution and hierarchical computation.

Tree shrews (*Tupaia belangeri*) provide a powerful system for extending our understanding of how biological object recognition depends on visual acuity and hierarchical processing. They are close relatives of primates (Janečka et al., 2007; Ni and Qiu, 2012) and share key features of primate vision, including a six-layered lateral geniculate nucleus, orientation maps in primary visual cortex, and an elaborated extrastriate cortex (Holdefer and Norton, 1995; Fitzpatrick, 1996; Van Hooser et al., 2013). Their cone-rich retina supports higher acuity than rodents (Petry et al., 1984; Grannonico et al., 2024), yet their peak cone density is lower than primates and they lack a foveal pit, resulting in blurrier optics and an acuity roughly an order of magnitude lower than in humans (Immel J.h.j., 1983; Müller and Peichl, 1989; Sajdak et al., 2019). Tree shrews are also dichromatic, possessing S- and L-cones but no M-cones, narrowing their color range relative to many primates (Jacobs and Neitz, 1986; Sajdak et al., 2019). Despite these constraints, cortical mapping studies reveal hierarchical progression beyond V1 and suggest a compact but organized network for visual form processing (Wong and Kaas, 2009; Lanfranchi et al., 2025). This combination of coarser front-end spatial resolution and structured cortical organization makes tree shrews ideal models to test whether invariant object recognition requires primate-level visual precision or can emerge within a reduced spatial sampling regime. Moreover, because tree shrews occupy a pivotal phylogenetic position near the base of the primate lineage, characterizing their visual capacities may shed light on evolutionary origins of primate high-level vision.

Here we tested whether tree shrews display behavioral signatures of transformation-tolerant object recognition and identified which visual features support this capacity. We modeled species-specific optics and retinal sampling by extending an image-computable model of the initial human visual encoding (Cottaris et al., 2019, 2020). This provides an estimate of what visual information is available for downstream processing (Zhang et al., 2022), allowing us to account for major differences in spatial acuity relative to primates and to assess whether their retinal filtering can preserve the relational structure among natural images. We then measured object recognition performance across changes in position, scale, and viewpoint and assessed generalization to novel exemplars. To assess robustness under more naturalistic viewing conditions, we also tested recognition of objects embedded within natural scene backgrounds. Finally, we compared behavioral performance with visual feature models and hierarchical neural network architectures to determine which types of visual information best predict recognition behavior. Together, these analyses reveal that tree shrew vision supports object-level recognition and generalization despite coarser front-end optics and cone sampling density.

## Results

### Modeling front-end visual information available to tree shrews

Tree shrews have roughly tenfold lower spatial acuity than humans, constrained by the optical properties and cone sampling density of their eyes, yet they are highly visual animals that rely on vision for navigation and foraging. To evaluate how these front-end differences constrain the visual input available to downstream visual areas for object recognition, we modeled the filtering imposed by the tree shrew eye to determine what visual information is preserved or lost in their initial visual encoding. This approach allowed us to estimate how much of the structure among natural objects, including distinctions relevant to visual categories and object recognition, survives optical limitations and could be transmitted to higher stages of the visual system.

We used the Image System Engineering Toolbox for Biology (ISETBio; (Cottaris et al., 2019, 2020) to simulate cone activations in response to natural images. The original model accounts for key front-end components of the human visual system, including stimulus display properties, ocular optics, and cone sampling density, enabling realistic estimation of the retinal image and mosaic of cone excitations under species-specific constraints. We adapted this model for tree shrews (“ISETTreeShrew”) to compare retinal-level visual representations across species. For both tree shrews and humans, we modeled retinal responses to a set of 92 grayscale images spanning faces, bodies, and inanimate objects (Kriegeskorte et al., 2008) at multiple image sizes (10°, 5°, 2.5°, and 1.25° diameter of visual angle [d.v.a.]), where image size corresponded to the length of the long axis for each stimulus. We then used a Bayesian reconstruction model (Zhang et al., 2022) to infer the retinal image representation from cone responses (Figure 1A).

**Figure 1:**
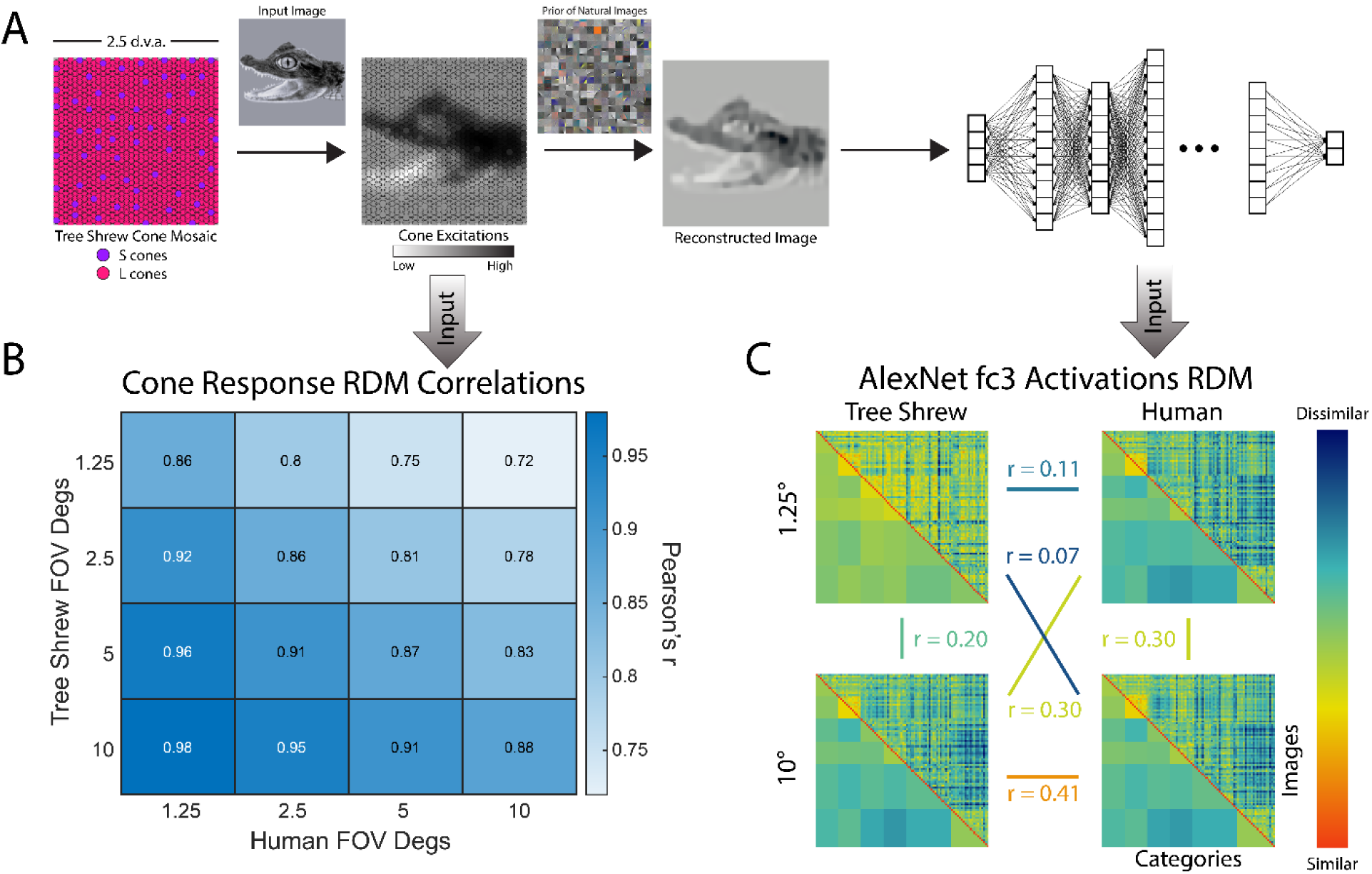
Tree shrew retinal filtering preserves object identity information at large image sizes. (A) Schematic of the modeling pipeline using the Image System Engineering Toolbox for Biology (ISETBio; Cottaris et al., 2019, 2020). The model incorporates species-specific front-end visual properties, including optical blur and cone mosaic density, to simulate cone activations in response to natural images. Bayesian reconstruction was then used to estimate image representations after retinal filtering. The sparse prior was built by fitting an exponential probability density function to weights derived by applying independent components analysis to a large dataset of natural images (Zhang et al., 2022). These reconstructed images were subsequently passed through a standard AlexNet model pretrained on ImageNet to examine representational geometry across hierarchical layers. Downward arrows indicate which data types are used for subsequent analyses. (B) Correlation matrix showing similarity (Pearson’s r) between tree shrew and human cone response RDMs across image sizes. Tree shrew and human retinal RDMs were most strongly correlated when object size was scaled for species-specific acuity (10° tree shrew ≈ 1.25° human). (C) RDMs computed by calculating distances between all 92 image pairs using activations from the final fully connected (fc3) layer of AlexNet. Upper triangles depict all image pair activations. Lower triangles depict average distance for each category pairing. Representative RDMs for tree shrew and human reconstructed images at 1.25° and 10° image sizes shown here. Cross-species correlations (Pearson’s r) show that representational geometry is most similar when images are scaled for acuity, indicating that tree shrew retinal filtering preserves structure sufficient for downstream object-level differentiation.

We first compared the visual information available for object recognition after retinal filtering by examining the pattern of distinctions among natural images in the estimated cone responses across tree shrews and humans. For each species, we constructed representational dissimilarity matrices (RDMs) by computing Euclidean distances between all image pairs based on modeled cone responses (Figure 1B). We then compared these matrices. This analysis captures how well the relationships among images are preserved despite optical blur and cone spatial sampling limitations. At larger stimulus sizes, the pattern of relationships among images in the tree shrew model closely matched that in the human model. The strongest correlation occurred between 10° images for tree shrews and 1.25° images for humans (Pearson’s r(4184) = 0.98, r > 97.5 percentile of null distribution from 1000 random permutations), consistent with the approximately tenfold difference in spatial acuity reported behaviorally (Petry et al., 1984). These results indicate that although tree shrew vision is spatially coarser, the retinal representation scaled to its acuity preserves a high degree of relational structure among natural images that could support category-level distinctions.

To assess how retinal filtering impacts the information structure available to downstream stages of visual processing, we reconstructed the modeled cone responses back into image space using a Bayesian reconstruction approach (Zhang et al., 2022) (Figure 1A). These reconstructions provide an estimate of the image information carried by the cone excitations, after passing through the optical and retinal sampling constraints of each species. The natural image prior incorporated into the Bayesian reconstruction approach sets these estimates in the context of natural viewing, and the results refer the visual representation back to the image domain allowing us to take advantage of image-based computational tools to make cross-species comparisons. We then passed the reconstructed images through a standard AlexNet model (Krizhevsky et al., 2012) pretrained on ImageNet (Deng et al., 2009). Using activations from each layer for all image sizes, we computed RDMs and assessed representational similarity across species (Figure 1A and C, Supplementary Figure 1A). Early convolutional layers primarily captured local texture statistics, whereas deeper layers reflected broader image relationships associated with object-level distinctions. For humans, object-level structure in the deepest layer was evident across all stimulus sizes. In contrast, tree shrew reconstructions showed size-dependent differences, particularly at the smallest image size. Although tree shrews and humans show some representational similarity in early layers for 2.5°, 5°, and 10° images, the strongest cross-species correlation occurred in the deepest full connected layer (fc3) for 10° images (Pearson’s r(4184) = 0.41, r > 97.5 percentile of null distribution from 1000 random permutations). These results suggest that, when scaled for acuity, retinal filtering in tree shrews preserves sufficient information to support object-level differentiation in downstream hierarchical models.

To characterize constraints on how object-level information is preserved within tree shrew vision, we next compared representational similarity across layers and image sizes within each species (Supplementary Figure 1B). Although tree shrews require larger images to achieve high correlations, both species show a similar relationship between image size, processing depth, and correlation with the largest image (10°) in the final layer (fc3). Together, these analyses suggest that preserving object-relevant structure in tree shrews requires larger stimulus sizes, effectively distributing object features over a broader extent of the visual field. Whether downstream visual areas in tree shrews can integrate information across this larger spatial range to support object recognition is therefore an empirical question, addressed in the following behavioral experiments.

### Tree shrews discriminate complex objects across identity-preserving transformations

To address whether tree shrews exhibit invariant object recognition, we tested their behavioral ability to discriminate between complex objects across transformations of position, scale, and viewpoint. Recognizing objects across such transformations requires integrating visual information over broad spatial extents, providing a stringent test of object recognition in a visual system lacking a fovea. We adapted a two-alternative forced-choice (2AFC) paradigm widely used in primate and rodent studies (Rajalingham et al., 2018; Vinken and Beeck, 2021; Kell et al., 2023), in which object identity and viewpoint typically vary across trials. When we initially applied this approach to tree shrews using a small stimulus set drawn from multiple object categories. On each trial a target object was presented centrally, followed by two choice objects that varied in position, scale, and viewpoint. One choice matched the target, and the other (the distractor) was from a different category. The animal indicated its choice via nose poke (Figure 2A) and was rewarded with juice for choosing the choice that matched the target. Image sizes subtended 10° of visual angle, the peak correspondence in category representations between tree shrews and human from the frontend optic modeling (Figure 1). In piloting experiments using 2D object photographs, tree shrews performed this task well when target identity varied across trials, but struggled when tested with 3D images. To reduce task difficulty while still probing transformation tolerance, we fixed the target identity to a single object category (a camel) and varied its translation, rotation, and scale. Tree shrews performed the task in a setup that allowed for free movement between a behavioral testing chamber and their home cage between trials, enabling high trial counts without head fixation.

**Figure 2:**
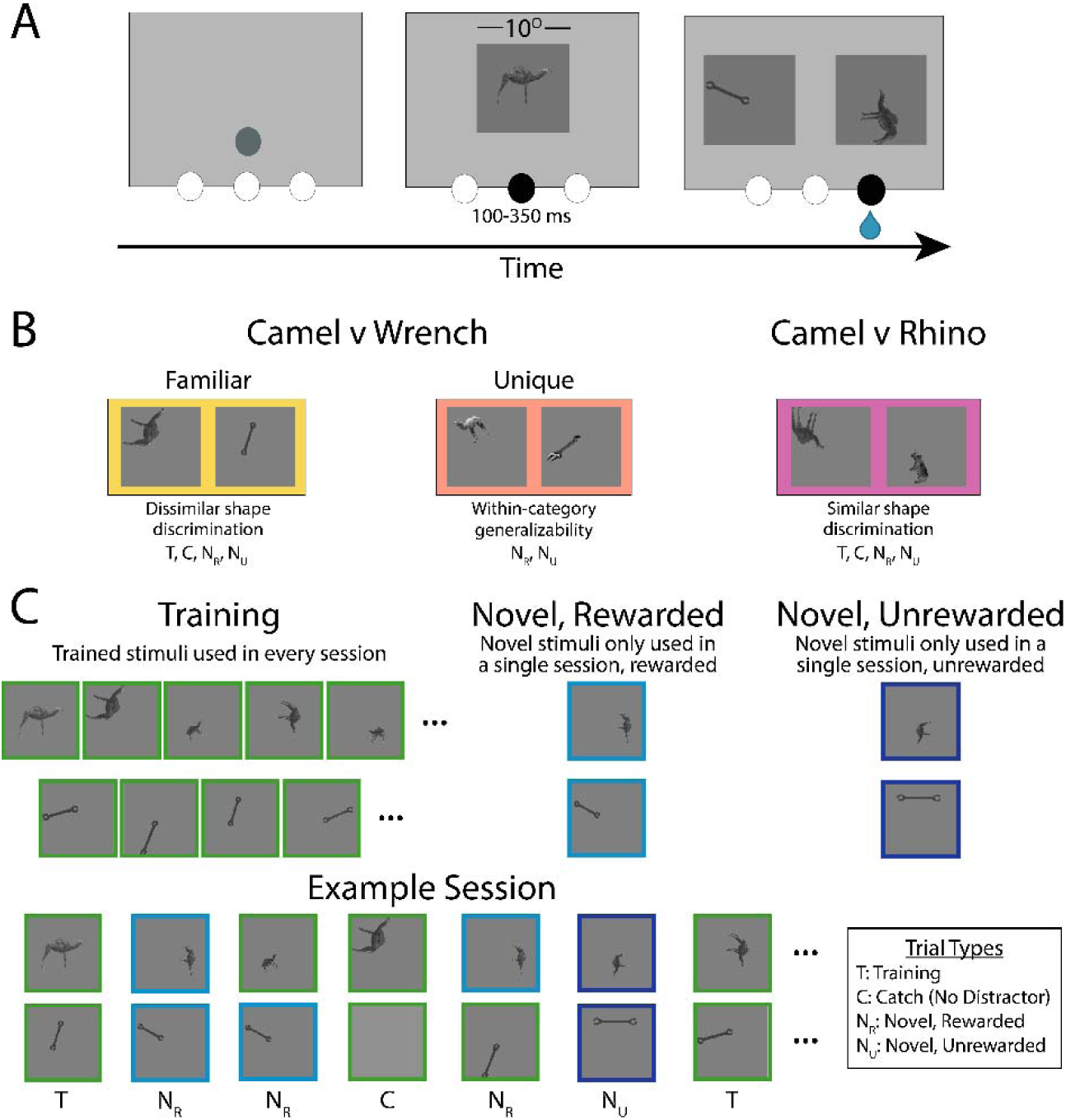
Schematic of task types and experimental design. (A) Behavioral task structure. Schematic illustrating the sequence of events within a single trial. Each trial began when the tree shrew initiated a central nose poke to reveal a reference image of the target (camel). After a brief delay, two choice images (10° of visual angle) that varied in position, scale and viewpoint appeared, one image depicting the target and the other a distractor (wrench or rhino). Correct choices triggered a juice reward. (B) Task variants. Schematics depicting the three task types used to manipulate object recognition difficulty. In the *camel versus wrench familiar* task, the same camel and wrench identities were shown across changes in position, rotation, and scale. In the *camel versus wrench* unique task, novel exemplars within each category were used to test generalization across untrained identities. In the *camel versus rhino* task, objects with more similar global shape were paired, creating the most challenging condition. (C) Stimulus sets for training and testing. Illustration of the stimulus types used in each session. Training stimuli were repeated across all sessions. *Novel rewarded* stimuli were new transformations of trained objects shown in a single session and rewarded for correct responses. *Novel unrewarded* stimuli were new transformations presented in a single session and never rewarded. Catch trials replaced the distractor with a mean gray square to assess upper-bound performance based on attention and motivation.

We studied three versions of the task designed to probe object recognition across increasing levels of difficulty (Figure 2B). In the *camel versus wrench familiar task,* tree shrews discriminated between two objects that were visually distinct in global shape across transformations of position, scale, and rotation. Both the camel target and wrench distractor identities used for testing were identical to those encountered during training, making this the easiest condition. The *camel versus wrench unique task* introduced novel exemplars from the same object categories that differed in fine structural details, such as a two-hump versus single-hump camel or a crescent versus box wrench. These images also differed in luminance and contrast relative to the trained images, allowing us to assess generalization across untrained identities within a category. Finally, the *camel versus rhino task* required discrimination between objects that were more similar in overall shape, representing the most difficult condition. Each animal was trained and tested sequentially on these task variants, advancing only after reaching stable performance and completing testing sessions on the preceding task.

For each task variant, tree shrews began training sessions with a single target-distractor pair. Additional stimuli were gradually introduced across sessions until performance plateaued, at which point the full training set contained 11 target images and 7 distractors per session spanning translation, rotation, and scale. After stable performance was reached, testing sessions were conducted to assess generalization to untrained transformations rather than reliance on memorized training examples (Figure 2C). Each testing session included two types of novel stimuli. *Novel rewarded* stimuli consisted of untrained transformations of the familiar target–distractor pair, presented only within a single session and rewarded for correct responses, providing a measure of generalization under reinforcement. *Novel unrewarded* stimuli were an additional pair of novel transformations presented only within a single session and never rewarded, providing a stricter test of generalization while preventing within-session learning. Only data from testing sessions were included in subsequent analyses.

All three animals successfully learned the *camel versus wrench familiar* task, two shrews completed the *camel versus wrench unique* task, and two completed the *camel versus rhino* task. Average individual performance across tasks ranged from 67–84%, slightly below performance on catch trials, which ranged from 84–94% (Figure 3A; Table 1). Even in the most difficult camel–rhino task variant, performance remained well above chance, demonstrating that tree shrews can recognize objects across substantial variation in position, rotation, and scale. Each session included several repetitions of every target–distractor pair, with a median of 864 trials per session (Supplementary Figure 2). This level of engagement is comparable to that reported for marmosets, a New World primate, performing similar paradigms (Kell et al., 2023) and provided sufficient repetitions to reliably estimate performance for individual target–distractor judgments.

**Figure 3:**
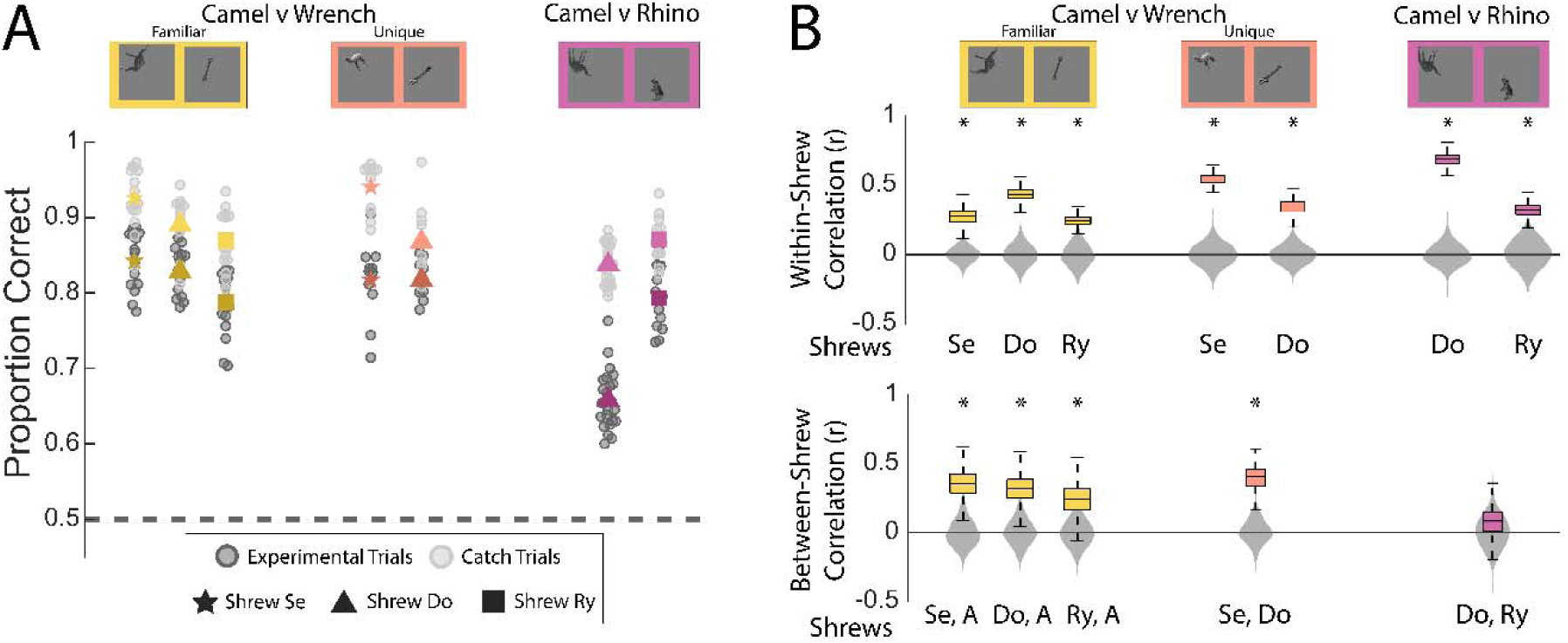
Tree shrews show consistent and shared object recognition behavior. (A) Object discrimination accuracy. Proportion correct for each tree shrew in the camel versus wrench and camel versus rhino tasks. Dark gray points indicate session averages for experimental trials, and light gray points show session averages for catch trials in which the distractor was replaced with a gray square. Each animal is represented by a distinct marker (Se = star, Do = triangle, Ry = square). Horizontal dashed line denotes chance performance (0.5). (B) Within- and between-shrew correlations. Box plots show distributions of correlation coefficients (r) for each task. The top row depicts within-shrew correlations calculated by randomly splitting trials for each target–distractor pair and correlating mean performance across splits. The bottom row shows between-shrew correlations of bootstrapped accuracy patterns for each pair of animals. Between-shrew correlations on the *camel versus wrench familiar* task were calculated by correlating an individual shrew’s performance with the average of the other two (A = average of other two shrews). Gray violin plots depict null distributions from random permutations. Asterisks denote correlations exceeding the 97.5 percentile of the null distribution.

**Table 1:**
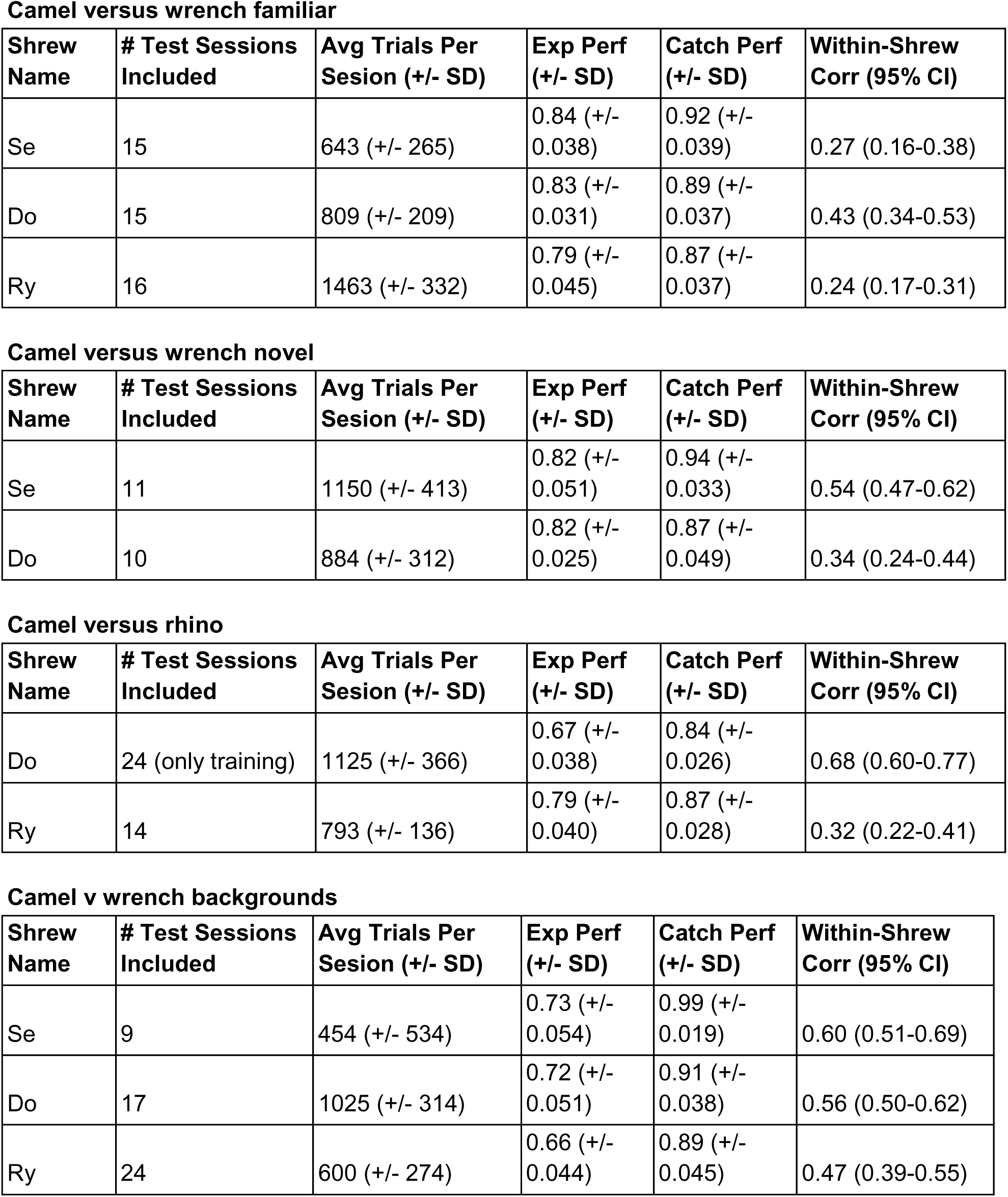
Summary statistics of task variations. Specific details of each task variation including shrews, session and trial counts, performance and consistency metrics.

To assess whether object recognition behavior reflected shared perceptual structure across individuals, we examined correlations in accuracy across stimulus pairs. For each shrew, we averaged performance across all target–distractor pairs and computed within-animal reliability by randomly splitting trials for each pair in half and correlating mean performance between splits. All three shrews showed significant within-animal correlations across tasks, indicating consistent and systematic response patterns across sessions (Pearson’s r > 97.5% of null distribution from permuted data; Figure 3B). We then compared performance patterns across animals to test whether similar perceptual structure guided their decisions. Between-animal correlations revealed significant similarity between shrews for both *camel versus wrench* tasks (Pearson’s r > 97.5% of null distribution from permuted target-distractor pairs). However, the correlation between shrews Ry and Do in the *camel versus rhino* task was not significant, likely reflecting reduced performance in this condition for shrew Do. Together, these results suggest that tree shrews rely on stable and largely shared perceptual strategies when discriminating objects, consistent with a common underlying visual architecture rather than arbitrary or idiosyncratic choices.

### Generalization to novel exemplars

A defining feature of object recognition is the ability to generalize to unfamiliar examples. To assess this capacity in tree shrews, we isolated trials from test sessions that contained either *novel rewarded* or *novel unrewarded* stimuli and analyzed performance across sessions (Figure 4A,B). Tree shrews generalized to novel rewarded stimuli in all task variants, performing well above chance despite never encountering these transformations during training (average shrew performance ranging from 76-84% across all tasks; Table 1). Performance on *novel unrewarded* stimuli was more variable (average shrew performance ranging from 59-77% across all tasks; Table 1), though accuracy remained above chance in most sessions.

**Figure 4:**
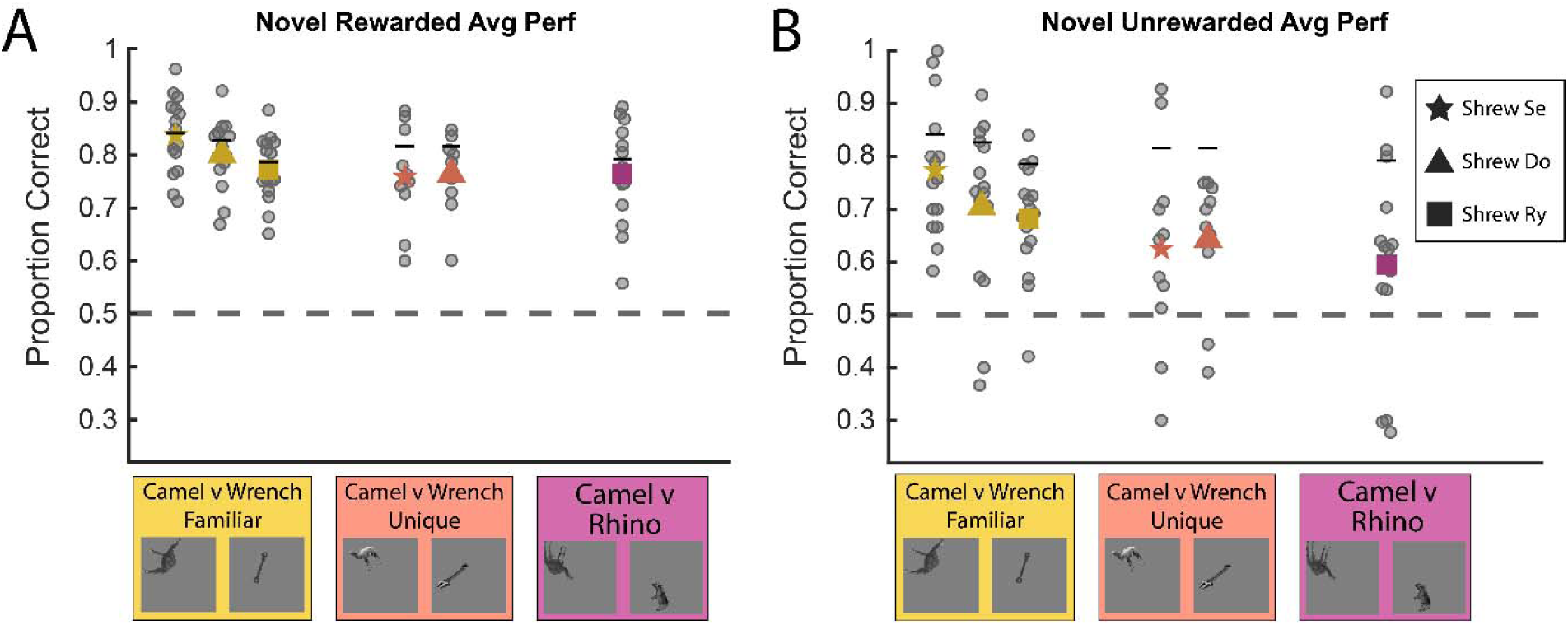
Tree shrew generalization performance for novel stimuli. (A) Novel rewarded trials. Proportion correct for each animal on camel versus wrench and camel versus rhino tasks using untrained transformations that were rewarded for correct responses. Gray points show individual session averages, and colored symbols indicate the mean performance for each shrew (Se = star, Do = triangle, Ry = square). Horizontal black lines indicate mean shrew experimental performance from Figure 3A. The horizontal dashed line denotes chance performance (0.5). (B) Novel unrewarded trials. Proportion correct for the same tasks using untrained transformations that were never rewarded, providing a measure of generalization without within-session learning. Plotting conventions match panel (A).

To determine whether performance on novel stimuli reflected rapid learning rather than generalization, we compared accuracy between the early and late halves of testing sessions. We observed no evidence of learning for either novel rewarded or novel unrewarded stimuli within sessions (Supplementary Figure 3), indicating that performance did not improve with repeated exposure. Together, these results show that tree shrews can generalize learned object identities to previously unseen exemplars, a core property of high-level object recognition.

### Influence of natural scene backgrounds

We next tested whether tree shrews could maintain object recognition when objects were embedded in natural scene backgrounds, introducing realistic visual clutter and segmentation demands. Animals were trained to discriminate camels from wrenches across six background images, each combined with multiple object transformations, yielding 18 unique target and distractor pairings. Average shrew accuracy on experimental trials ranged between 66-73%, indicating that tree shrews could perform the task despite increased visual complexity (Figure 5A). Performance on catch trials, in which the distractor was replaced with a gray square, remained near ceiling (ranging between 89-99%), likely reflecting the salient contrast between object images and a blank alternative. These results show that tree shrews can successfully recognize objects even when they are embedded in complex, naturalistic scenes, although performance is reduced compared with simpler backgrounds.

**Figure 5:**
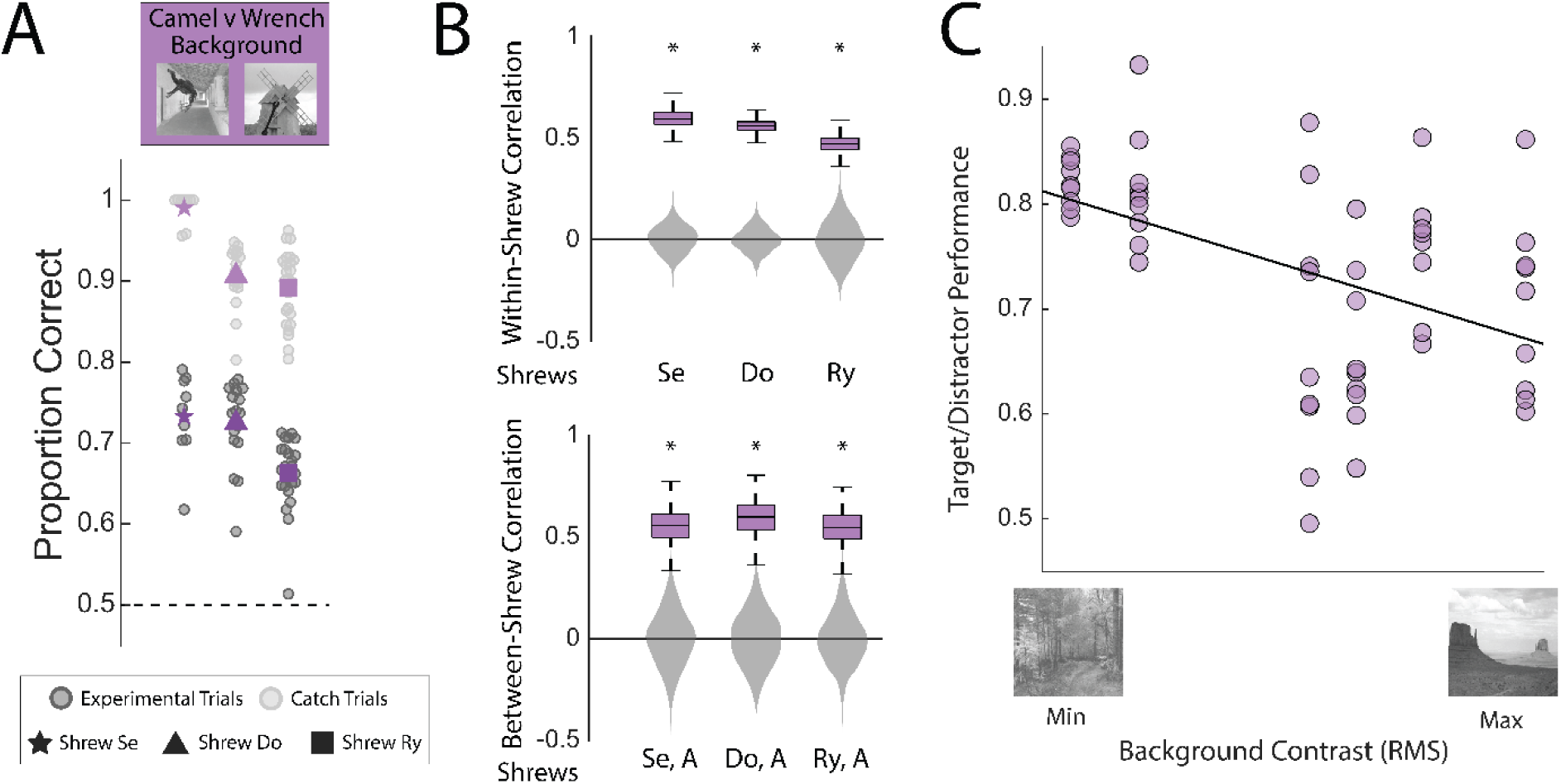
Tree shrew object recognition in natural scene backgrounds. (A) Behavioral performance on cluttered scenes. Proportion correct for each animal on the camel versus wrench task with natural scene backgrounds. Gray points show individual session averages for experimental (dark) and catch (light) trials. Each animal is represented by a distinct marker (Se = star, Do = triangle, Ry = square). The horizontal dashed line denotes chance performance (0.5). (B) Consistency of response patterns. The top row depicts within-shrew correlations calculated by randomly splitting trials for each target–distractor pair and correlating mean performance across splits. The bottom row shows between-shrew correlations of bootstrapped accuracy patterns calculated by correlating an individual shrew’s performance with the average of the other two (A = average of other two shrews). Gray violin plots depict null distributions from random permutations. Asterisks indicate correlations exceeding the 97.5 percentile of the null distribution. (C) Influence of background contrast on accuracy. Scatterplot showing the relationship between background contrast (root mean square; RMS) and target–distractor accuracy. Individual points represent average performance on each target-distractor pair for shared background presentations. The line indicates the best-fit linear regression. Example backgrounds with minimal and maximal contrast are shown below the x-axis. Pearson’s r(51) = - 0.46, p = 0.00049.

To determine whether performance patterns were systematic, we again computed within-animal split-half reliability and between-animal correlations. All within-animal correlations were significant, as were between-animal correlations for all pairs (Pearson’s r > 97.5% of null distribution from permuted data; Figure 5B), indicating consistent recognition structure across individuals. However, accuracy patterns for objects embedded in scenes did not correlate with performance on the same objects presented without backgrounds (Pearson’s r(7) = -0.09, p = 0.82; Supplementary Figure 4), suggesting that scene context alters the effective information guiding recognition. To evaluate one low-level feature that may impact performance, we examined background contrast and found that higher contrast was significantly associated with lower accuracy across all stimulus pairs (Pearson’s r(51) = -0.46, p = 0.00049; Figure 5C). These results are consistent with prior findings in macaques showing that object recognition performance varies with foreground-background contrast (Rajalingham et al., 2018). Together, these results show that while tree shrews can recognize objects in scenes, performance is modulated by background image statistics, highlighting the importance of accounting for scene-driven interference when assessing object recognition under naturalistic contexts.

### Modeling visual feature complexity underlying object recognition behavior

Having established that tree shrews can recognize objects across transformations and within natural scenes, we next asked what kinds of visual information support this behavior. Specifically, we tested whether performance patterns could be explained by low-level image properties or instead reflect sensitivity to more complex representations that emerge through hierarchical processing. To focus on shared perceptual structure, we analyzed average performance across all three animals from the camel versus wrench familiar task. Analyses were restricted to training images presented during test sessions (84 unique target-distractor pairs), which had the highest trial counts and provided the most reliable estimates of behavioral performance.

Previous work in rodents and primates has shown that object discrimination performance is strongly related to object size (Rajalingham et al., 2018; Schnell et al., 2023a). Consistent with these findings, tree shrew performance was significantly correlated with the relative target-distractor size difference (r(82) = 0.388, p = 0.00026; Supplementary Figure 5). Accuracy patterns across target–distractor size combinations suggested that larger objects enhance perceptual salience and the resolution of object features relevant for discrimination. Importantly, performance remained above chance even when distractors were larger than targets (Supplementary Figure 5), demonstrating that the shrews weren’t simply choosing the largest image.

To explore what additional visual information beyond object size could support object discrimination, we constructed a set of candidate visual feature representations spanning multiple levels of complexity. These representations sampled various aspects of image structure, from raw pixel similarity to more abstract shape-based descriptions. To test whether performance could be explained by image-level similarity while accounting for the fact that tree shrews were freely moving and could compensate for differences in object position or scale, we created original images (“*object*”), translation-normalized (“*object centered*”), scale-normalized (“*object scaled*”), and jointly-normalized images (“*object centered and scaled*”), in which position and/or size were matched to the reference camel. All images were reconstructed using the ISETTreeShrew model subtending 10° of visual angle.

We next constructed feature-based representations from ISETTreeShrew reconstructed images designed to capture visual properties beyond raw pixel similarity. Lower-level feature models included a Gabor filter representation capturing orientation and spatial frequency content, as well as two texture-based models. One model synthesized texture across the full image (“*texture*”), whereas a second preserved coarse object position and size information by computing texture within object boundaries (“*texture-in-place*”). To test whether global shape contributed to performance, we generated binarized object silhouettes (“*binary object*”) and medial-axis skeleton representations (“*skeleton*”), which emphasize object outline and topology while minimizing local texture cues. To isolate shape information from size, we additionally created size-normalized variants (“*binary object, scaled*” and “*skeleton scaled*”). Finally, we included a *saliency* model trained on human eye fixations (Jiang et al., 2015) to assess whether predicted visual attention could account for performance patterns. For all non-pixel representations, model outputs were passed through shallow convolutional networks trained to discriminate targets from distractors. This reduced sensitivity to precise spatial alignment while preserving feature content and produced representational distances that enabled comparisons across feature model types (See “Modeling of tree shrew behavior” in Methods).

Because these pixel- and feature-based models capture partially overlapping visual information, they do not constitute independent descriptions of the stimuli. We therefore computed pairwise correlations between target-distractor distance patterns across all models and performed hierarchical clustering on this correlation matrix (Supplementary Figure 6). This analysis revealed two main clusters and a set of unclustered models. Cluster 1 comprised representations in which object size was explicitly normalized across stimuli (including scaled pixel and scaled shape representations), whereas Cluster 2 consisted of shape-based representations that preserved original object size information. Unclustered models primarily reflected lower-level feature statistics such as textures, orientation, or saliency (see dendrogram in Figure 6A). We then correlated the pairwise distances between target-distractor pairs for each model representation with patterns of tree shrew behavioral performance on the corresponding stimuli to estimate what information tree shrews may be using to make their decisions (Figure 6A). None of the unclustered models were significantly correlated with behavioral performance across target-distractor pairs (texture r = 0.011, 95% CI = [-0.199, 0.193]; Gabor r = -0.001, 95% CI = [-0.163, 0.183]; saliency r = -0.139, 95% CI = [-0.310, 0.055]; texture-in-place r = 0.188, 95% CI = [-0.009, 0.395]). Within Cluster 2, shape-based representations that preserved object size showed significant correlations with behavior (skeleton r = 0.263, 95% CI = [0.158, 0.389]; binary r = 0.35, 95% CI = [0.20, 0.51]), whereas pixel-based representations showed weaker or non-significant relationships (object r = 0.10, 95% CI = [-0.10, 0.36]; object centered r = 0.18, 95% CI = [-0.03, 0.40]). In contrast, Cluster 1 pixel representations, in which size information was explicitly removed, did not positively correlate with behavior (object scaled r = -0.258, 95% CI = [-0.402, -0.103]; object centered and scaled r = -0.160, 95% CI = [-0.311, -0.011]). Similarly, removing size information from shape representations eliminated their correspondence with behavior (skeleton scaled r = -0.007, 95% CI = [-0.22, 0.17]; binary scaled r = -0.12, 95% CI = [-0.30, 0.06]). Together, these results indicate that the correlation between shape-based models and behavior depends on preserving relative target–distractor size differences. Removing size information abolishes this relationship, suggesting that pairwise size contrast is a major contributor to discrimination performance. This is consistent with our earlier analyses (Supplementary Figure 5), which show that behavioral accuracy tracks relative size differences, although performance remains above chance even when size favors the distractor.

**Figure 6:**
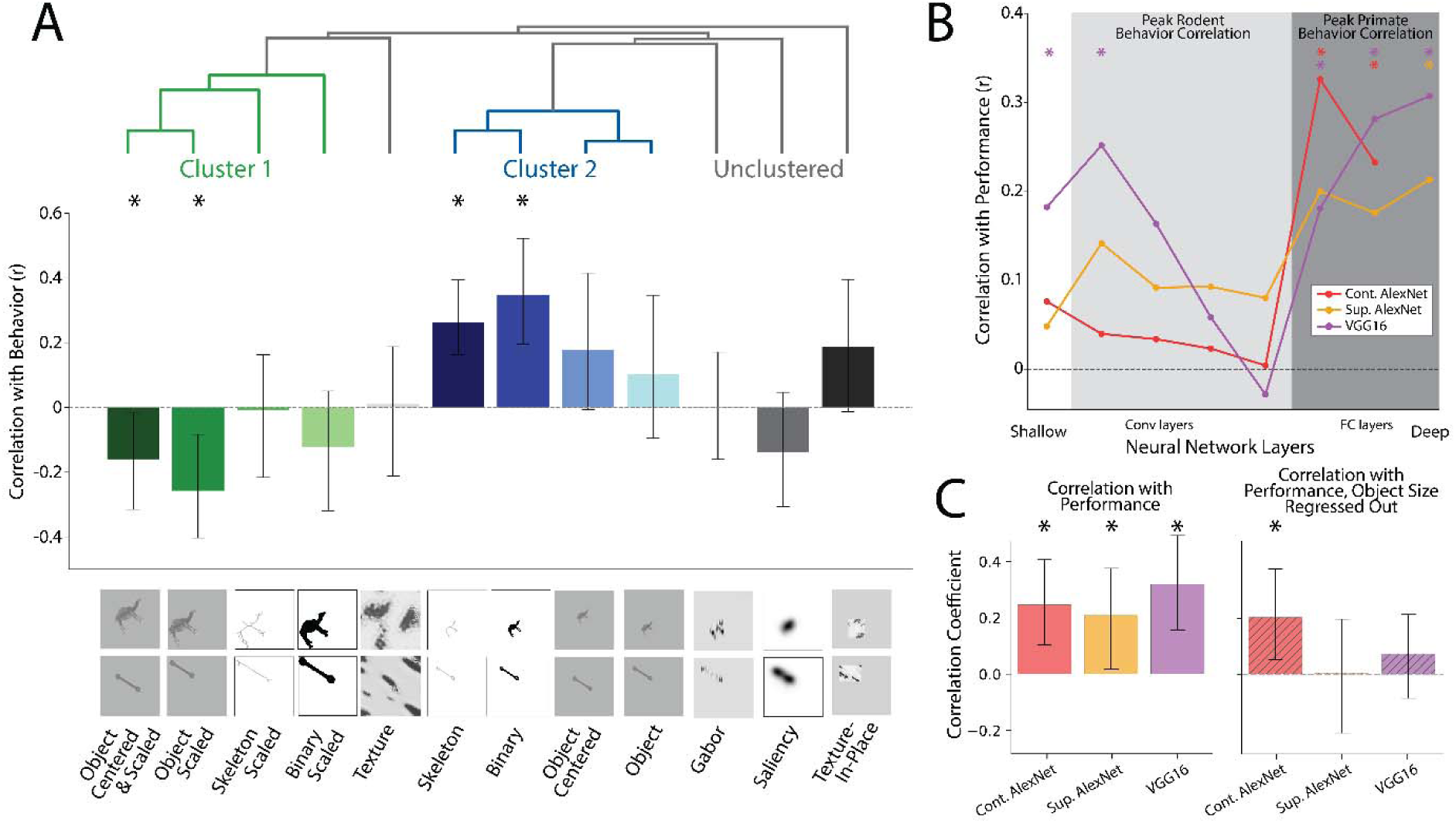
Modeling visual features underlying tree shrew object recognition behavior. (A) Correlation between tree shrew behavioral performance and representational distances derived from different visual feature models. Dendrogram depicts hierarchical clustering of model correlation matrix (Supplementary Figure 6). Bar plots show correlations between average behavioral performance and representational distances between targets and distractors from different model classes. Example representations for each model type are shown below. Error bars represent 95% confidence intervals from 1000 bootstrapped samples. (B) Behavioral correspondence across model hierarchies. Correlation between behavioral performance and representational distances computed from successive layers of supervised AlexNet, contrastive AlexNet, and VGG16 models applied to ISETTreeShrew-reconstructed images. Shaded regions indicate the approximate layer range showing peak correspondence with mouse and primate behavioral data in previous studies (Nayebi et al., 2023; Vinken and Beeck, 2021; Hong et al., 2016; Kell et al., 2023). Asterisks (*) indicate correlations exceeding 97.5 percentile of null distribution from 1000 random permutations. (C) Correlation between behavioral performance and representational distances from final layer of each neural network. Striped bars show correlations after regressing out object size differences. Error bars represent 95% confidence intervals from 1000 bootstrapped samples.

Whereas these explicit feature models were designed to test the contribution of specific visual features to behavior, we next used deep neural network models to assess how correspondence with tree shrew performance varies across levels of hierarchical visual processing. We analyzed three models pretrained on ImageNet (Deng et al., 2009): a classic supervised AlexNet model (Krizhevsky et al., 2012), a contrastive AlexNet model used as a closer comparison to rodent visual systems (Nayebi et al., 2023), and VGG16 (Simonyan and Zisserman, 2015). For each network, we extracted activation distances from successive convolutional and fully connected layers in response to the ISETTreeShrew-reconstructed training images and correlated these distances with tree shrew performance (Figure 6B). All three networks showed peak correspondence in the final layers, with VGG16 showing an additional peak in its earliest layers. This depth-dependent pattern of correspondence with tree shrew behavior (Figure 6B) is more aligned with primate object recognition, in which behavioral correspondence is strongest in deep layers (Yamins et al., 2014; Hong et al., 2016; Kell et al., 2023), and contrasts with rodent vision, where correspondence peaks at intermediate layers (Vinken and Beeck, 2021; Nayebi et al., 2023; Schnell et al., 2023a).

Given the association between relative target-distractor size differences and behavioral performance, we next examined neural network-behavior correlations after regressing out size effects (Figure 6C). After size regression, correlations for both supervised models were eliminated, whereas the contrastive model retained a significant correlation with behavior. These results indicate that while relative object size contributes significantly to behavioral performance, higher-level representations can capture additional structure beyond size alone. Contrastive, self-supervised learning explicitly promotes similarity across augmented views of the same image while separating representations of different objects (Chen et al., 2020; Tian et al., 2020; Ge et al., 2022). By aligning representations across transformations that include changes in scale, position, and other low-level image properties, contrastive objectives reduce dependence on image-level cues that covary with object size and instead emphasize more transformation-tolerant shape structure. This property may account for the contrastive model’s closer correspondence with tree shrew behavior after controlling for size differences.

## Discussion

Tree shrews recognized objects across changes in position, scale, and viewpoint, generalized to novel exemplars, and identified objects embedded in natural scenes. Behavioral patterns were generally reliable across sessions within animals and correlated across animals, suggesting that the shrews relied on consistent visual cues rather than idiosyncratic associations specific to an individual shrew’s learning. Front-end optical modeling showed that scaling images to account for the coarser spatial sensitivity of tree shrew vision (Petry et al. 1984) preserves sufficient relational structure at the retinal level to support these object-level distinctions. Despite this scaling, relative object size in the image accounts for substantial structure in both primate and tree shrew behavioral performance (Rajalingham et al., 2018) and in late-stage hierarchical neural network representations, indicating that increased object size inherently retains perceptual clarity rather than influencing behavior optical limitations. Importantly, object size did not fully account for behavioral structure, suggesting that additional visual information contributes to tree shrew object recognition. Together, these results demonstrate that the tree shrew visual system supports transformation-tolerant object recognition through hierarchical visual processing despite coarser front-end sampling. This establishes tree shrews as a valuable comparative model for investigating the neural basis and evolutionary origins of high-level vision.

### Object recognition and generalization in tree shrews

Most prior behavioral studies in tree shrews have focused on low-level visual discrimination. Early work established that tree shrews have higher spatial acuity and contrast sensitivity than rodents, consistent with their cone-dominated retina and diurnal lifestyle (Petry et al., 1984). However, the behavioral paradigms used have largely involved simple pattern or grating discrimination (Khani and Rainer, 2012; Li et al., 2022; Schumacher et al., 2022), which do not resolve whether tree shrews can recognize complex objects or generalize object identity across changes in viewing conditions. More recent work has examined figure-ground segmentation, a key intermediate visual process that delineates objects from backgrounds, using gratings and noise textures, showing that both tree shrews and rodents rely heavily on local cues when distinguishing textures (Luongo et al., 2023). The current results extend this literature by demonstrating that tree shrews can integrate visual information across position, scale, and viewpoint to recognize objects. Recognition when objects were embedded in natural scenes further demonstrates tolerance to visual clutter and contextual variation. Further, generalization to novel exemplars indicates that object representations are not tied to individual learned views. Behavioral performance aligned with late-stage hierarchical representations from a contrastively trained neural network model, consistent with representations that emphasize invariant similarity across changes in image scale (Chen et al., 2020; Tian et al., 2020; Ge et al., 2022). Together, these findings indicate that tree shrews engage intermediate and high-level visual processing mechanisms essential for robust object recognition across changes in appearance, providing a behavioral foundation for linking object recognition performance to specific cortical computations in a non-primate visual system.

### Tree shrews in a comparative context

The behavioral and modeling results suggest that the tree shrew visual system supports computations necessary for transformation-tolerant object recognition within a hierarchically organized cortical network. Situating tree shrew vision within the broader landscape of mammalian visual systems provides insight into the evolutionary expansion of visual processing hierarchies observed in primates.

The rodent visual system has been described as a relatively shallow and resource-limited network, relying on broadly tuned representations rather than deeply specialized feature hierarchies (Nayebi et al., 2023). Correspondingly, rodent object recognition performance is best captured by early to intermediate stages of hierarchical models, with limited additional explanatory power from deeper network layers (Vinken and Beeck, 2021; Nayebi et al., 2023). Rodent generalization abilities often break down under substantial variation in viewpoint or visual context. In contrast, tree shrews generalized across novel exemplars and cluttered scenes, and their behavioral patterns were best predicted by late-stage hierarchical representations, a profile more similar to primates than rodents. This difference suggests that object recognition in tree shrews relies on integrative computations that extend beyond those typically sufficient to account for rodent performance. This distinction is consistent with prior cross-species findings showing that tree shrews remain visually guided as task demands increase, whereas rodents increasingly disengage from visual input and adopt non-visual strategies (Mustafar et al., 2018).

The primate visual system comprises expanded, serially organized hierarchies that support robust category-invariant recognition (Hung et al., 2005; Yamins et al., 2014; Kell et al., 2023). Tree shrews share several key architectural features with primates, including dense cone mosaics, structured orientation maps, and an elaborated extrastriate cortex, which are absent in rodents (Petry and Bickford, 2019). The present findings extend this anatomical and physiological continuity to behavior, showing that these shared features are accompanied by object recognition strategies that rely on integrated, hierarchical processing. Together, this evidence positions tree shrews between rodents and primates, with behavioral signatures more consistent with deeper hierarchical computation than typically observed in rodents.

Comparative work in birds suggests that the capacity for object recognition extends beyond the mammalian lineage. Pigeons can learn to categorize complex objects and generalize to new exemplars, yet their transformation tolerance depends strongly on exposure to multiple views during training (Wasserman et al., 1996; Peissig et al., 2002; Soto and Wasserman, 2014). Tree shrews, by contrast, generalized flexibly to novel object identities without requiring broad exemplar experience during training, suggesting that this capacity reflects a more robust representational substrate than has been documented in birds. The extent to which this generalization scales across more challenging stimulus conditions, including greater dissimilarity between trained and novel exemplars and more similar competing categories, remains to be determined. Such comparisons across diverse vertebrate lineages will clarify the degree to which transformation-tolerant object recognition reflects shared computational solutions shaped by convergent pressures on visual systems, and how these capacities relate to specific features of cortical organization.

### Neural architecture underlying object recognition behavior

The behavioral findings raise a central question about the cortical architecture supporting tree shrew visual behavior. Tolerance to changes in position, scale, and viewpoint implies the existence of circuits that integrate across spatial transformations to support stable representations despite varying retinal inputs. Such integration extends beyond early retinotopic coding and is a hallmark of hierarchical visual processing in primates (Riesenhuber and Poggio, 1999; Hong et al., 2016). Visual cortical maps are highly conserved across primates (Arcaro and Kastner, 2015; Meyer et al., 2025), suggesting that hierarchical organization can be maintained across substantial evolutionary expansion. One possibility is that tree shrews possess a similarly structured visual hierarchy within a smaller cortical substrate. Alternatively, they may achieve comparable functional outcomes through a more condensed hierarchical organization (Nayebi et al., 2023).

Most electrophysiological studies in tree shrew visual cortex have focused on primary visual cortex (Van Hooser et al., 2013; Schumacher et al., 2022), but recent work has begun to characterize processing beyond V1. (Poirot et al., 2016) reported increased selectivity for grating structures from V1 to V2, with V2 neurons showing enhanced responses to polar gratings at high spatial frequencies, consistent with an emerging sensitivity to curved contours and scene boundaries. (Lanfranchi et al., 2025) further showed progressive increases in receptive field size, response latency, and selectivity for complex textures across V2 and higher-order visual areas, indicating the presence of increasingly integrative visual representations along the cortical hierarchy. Notably, neural responses in tree shrew V2 are best explained by intermediate and late layers of deep neural networks, suggesting that object-related processing depends on higher-order feature integration within a relatively compact cortical network (Lanfranchi et al., 2025). Such a compact architecture may support computations that, in primates, are distributed across multiple ventral stream areas. The sinusoidal transformation of retinotopy in V2 (Sedigh-Sarvestani et al., 2021) provides one potential mechanism, allowing spatially distributed visual inputs to converge while preserving local continuity and potentially facilitating integration across extended regions of visual space earlier in the processing hierarchy than in primates. This organization could enable hierarchical feature integration sufficient for transformation-tolerant object recognition despite a smaller cortical substrate. Together, the behavioral and physiological evidence suggests that core hierarchical computations associated with object recognition in primates are present in tree shrews, potentially implemented within a more compact cortical framework. Direct recordings from higher-order visual areas will be necessary to test these hypotheses more directly.

### Constraints of task design on cross-species comparisons

The comparative framing above assumes that behavioral tasks engage comparable computations across species. Even subtle differences in task design, however, can shift which aspects of visual processing are engaged (Kay et al., 2023). In object recognition, one important dimension is whether objects are presented in isolation or embedded within scenes. Embedding objects within scenes provides a more stringent and ecologically relevant test of recognition, but it also introduces additional computational demands. Performance in such cluttered conditions depends not only on encoding object-specific features but also on engaging mechanisms of figure–ground segregation, border assignment, and contextual integration. These mechanisms may differ across species, as may where they operate along stages of the ventral visual hierarchy (Luongo et al., 2023). Our findings further emphasize that recognition accuracy therefore reflects a combination of object-level encoding and figure–ground segregation that is required for scene processing.

Comparative evidence supports this interpretation. In primates, ventral stream representations preserve object identity despite visual clutter (Rust and DiCarlo, 2010), whereas in rodents, variation in scene structure often disrupts discrimination (Zoccolan et al., 2009). Tree shrews appear to fall close to primates on this spectrum, recognizing object identity across transformations when objects are embedded in scenes while remaining sensitive to variation in scene content (Rajalingham et al., 2018). This profile is consistent with a representational system that supports object-level structure without full invariance to scene content. Direct comparisons of identical objects across scene contexts, including manipulations that vary scene structure independently of object identity, will be necessary to isolate object-specific computations from scene-dependent influences and to more precisely position tree shrew vision along the continuum of contextual processing across species.

Task design also shapes performance through training structure and motivation (Kay et al., 2023). Cross-species work shows that stable object recognition performance often requires adjusting task complexity, including the number of objects and response contingencies, across primate species (Kell et al., 2023). Accordingly, we adopted a simplified design with a constant target identity and a limited object set, while maintaining motivation through interleaving familiar and test trials. This was particularly important in a freely-moving, in-cage paradigm, where tree shrews could disengage if task demands were too high. These design considerations highlight the importance of aligning task structure with species-specific behavioral capacities when interpreting cross-species differences in object recognition. Future refinements, including more naturalistic response demands, increased choice alternatives, or systematic manipulation of reward contingencies, may improve sensitivity while preserving comparability across species.

### Future directions

The present findings open several avenues for studying how tree shrews represent and generalize visual information. Recognition of new exemplars, such as single- versus double-humped camels and box versus crescent wrenches, indicates generalization beyond memorized object views. Future studies can expand the object space beyond a single target–distractor axis to include families of related objects that vary along graded diagnostic features. Such designs would permit direct tests of learning categorical representations and quantifying generalization across feature similarity distances. Systematic manipulation of object discriminability and introduction of novel exemplars that preserve category structure while varying surface details would further clarify whether tree shrew object representations support flexible category formation.

The behavioral results also generate concrete predictions about underlying neural mechanisms. Our network model comparisons suggest that recognition depends on intermediate and late stages of hierarchical processing that emphasize global shape and tolerance to transformations. Converging physiological evidence suggests that such computations may arise within a compact cortical hierarchy spanning V2 and nearby regions (Lanfranchi et al., 2025), but the spatial distribution and functional specialization of these representations remain unresolved. Combining object recognition behavior with neural recordings or imaging will be essential for determining whether hierarchical codes are localized to specific extrastriate areas or distributed across broader cortical networks. Establishing this link will clarify how cortical architecture constrains representational capacity and how hierarchical processing scales across species.

## Conclusion

Together, these findings demonstrate that tree shrews likely achieve visual object recognition through structured, hierarchical processing that extends beyond early sensory encoding. Their behavior reflects intermediate stages of visual computation that bridge rodent and primate vision, illustrating how tolerance to object transformations and visual context can emerge across species. By integrating behavioral, computational, and anatomical evidence, this work establishes the tree shrew as a powerful comparative model for studying the neural and evolutionary origins of object recognition. Future studies combining electrophysiology, imaging, and causal manipulation will be essential for determining how specific cortical circuits implement these computations and how they scale with evolutionary expansion. Elucidating these mechanisms in tree shrews will help identify general principles by which visual systems construct stable representations of objects across changing sensory inputs.

## Methods

### Animals

Three adult male tree shrews (*Tupaia belangeri*), aged 4–7 years, participated in this study. Animals were housed individually in a temperature- and humidity-controlled room on a 12-hour light/dark cycle. Food and water were available ad libitum, except during behavioral testing sessions when water bottles were removed. All experimental procedures complied with the National Institutes of Health Guide for the Care and Use of Laboratory Animals and were approved by the University of Pennsylvania Institutional Animal Care and Use Committee (IACUC).

### Visual Stimuli

Two stimulus sets were used: one for front-end modeling of retinal input and one for behavioral testing.

#### Stimuli for front-end modeling

We used 92 grayscale natural images spanning faces, bodies, and inanimate objects (Kriegeskorte et al., 2008). Each image was modeled with ISETBio and ISETTreeShrew and simulated at image sizes subtending 1.25°, 2.5°, 5°, and 10° of visual angle (d.v.a.). Images were converted to grayscale, then cone activations and image reconstructions were computed at the specified sizes. Mean luminance and root-mean-square (RMS) contrast were matched across sizes prior to modeling. These stimuli were used to estimate how optical blur and retinal sampling affect representational geometry in tree shrews and humans, and to generate species-specific reconstructions used for downstream analyses.

#### Stimuli for behavioral testing

Stimuli were drawn from a 3D object dataset developed by (Rajalingham et al., 2015) and later adapted by (Kell et al., 2023). Three object categories were used: camel, wrench, and rhino. All images were grayscale renderings that varied systematically in 3D rotation, translation, and scale to test generalization across identity-preserving transformations.

For the camel versus wrench tasks, the stimulus set included twelve camel images and seven wrench images derived from a single 3D model for each category, as well as additional exemplars used to test generalization across object identities, such as single- versus double-hump camels and box versus crescent wrenches. For the camel versus rhino task, eleven camel images and seven rhino images were used, varying across the same transformation parameters.

To evaluate recognition under naturalistic conditions, a subset of camel and wrench images was overlaid on natural scene backgrounds. Each object transformation (three per category) was combined with six backgrounds, yielding eighteen unique target and eighteen unique distractor images. Transformation–background pairings were rotated across sessions until all combinations were shown.

### ISETBio Tree Shrew Model

The ISETBio model of the human initial visual encoding was customized to match known properties of the tree shrew visual system. Below we outline how the key ISETBio objects were customized to allow computation of the tree shrew retinal image and cone excitations. In our simulations, both species’ models used an integration time of 1 ms and mean luminance of 20 cd/m^2^.

#### Lens

A custom ISETBio lens object was generated using the combined absorbance spectra of the tree shrew cornea and lens as measured by the study of (Petry and Harosi, 1990) by digitizing the data in Figure 5 of that study.

#### Physiological optics

Custom physiological optics models were derived based on a 4.35 mm post-nodal distance (Callahan and Petry, 2000), and the monochromatic wavefront aberration measurements in 11 tree shrews from the study of Sajdak et al., 2019. The wavefront measurements from that study were provided to us in tabular form by Austin Roorda (personal communication). From these data, we derived polychromatic point spread functions (PSFs) that accounted for chromatic aberration. We did this by adding to the defocus Zernike coefficient reported by Sajdak et al., 2019 a wavelength-dependent defocus amount which was computed from a single refracting surface tree shrew model eye, as derived in Sajdak et al., 2019.

#### Cone photoreceptors

Estimates of tree shrew cone absorptance spectra were generated for tree shrew L- and S-cones starting with the absorbance spectra measured by Petry and Harosi, 1990, by digitizing Figures 2 and 3 of their work. There are no M-cones in the tree shrew retina (Müller and Peichl, 1989; Petry and Harosi, 1990). Photopigment axial density was estimated for both L- and S-cones by assuming the human L-cone specific density and estimates of tree shrew cone outer segment lengths (Samorajski et al., 1966).

#### Cone mosaic

The tree-shrew retina does not contain a macular specialization or a macular pigment (Müller and Peichl, 1989; Sajdak et al., 2019), so the macular pigment object in the ISETBio cone mosaic model was set with an optical density of 0. Since the topography of cones in the tree shrew retina does not exhibit a density variation along the horizontal meridian, and only has a small variation along the vertical meridian (Sajdak et al., 2019), a custom ISETBio cone mosaic model for the tree shrew was generated based on a regular, eccentricity-independent hegaxonal structure. The retinal magnification factor was set to 76 microns/deg, corresponding to the 4.35 mm posterior nodal distance (Callahan and Petry, 2000). Cone spacing was set to 6.0 microns and cone aperture was set to 5.5 microns, which resulted in a cone mosaic model with a density of 32,300 cones/mm^2^. This value agrees with the maximum cone density ranges reported by Müller and Peichl, 1989 (around 32,000), and by Sajdak et al., 2019 (33,366 to 24,489). The relative density of L- and S-cones was set to 93/7, based on the 4% - 10% range in the relative proportion of blue-sensitive cones reported by Petry and Harosi, 1990. The position of S-cones in the cone mosaic lattice was adjusted to be semi-regular following the observations of Petry and Harosi, 1990 (their Figure 9).

Supplementary Figure 7 compares the ISETBio models of the cone mosaic, physiological optics, and longitudinal chromatic aberration (LCA) of tree shrews with those of humans, both for typical subjects (Polans et al., 2015; Sajdak et al., 2019). Note that retinal cone contrast in tree shrews is attenuated significantly more than in humans, due to all components of the front end: larger cones, more diffuse optics, and a stronger LCA across the visible spectrum.

#### Image reconstructions

Cone activations generated by this model were used to reconstruct how images would appear to tree shrews using the Bayesian inference framework (Zhang et al., 2022). Reconstructions were generated for both humans and tree shrews at image sizes subtending 1.25°, 2.5°, 5°, and 10° d.v.a. To minimize border artifacts, cone mosaics were expanded by 20%, with proportional image padding before cropping reconstructions back to their original size. To reduce computational load, render matrices and reconstructions were computed in a piecewise manner by reconstructing overlapping blocks across the visual field then merging afterwards. Reconstruction parameters were 4,000 iterations with λ = 0.001 for humans and λ = 1,000 for tree shrews. See Zhang et al., 2022 for description of the image prior used in the reconstruction and the meaning of parameter λ. We applied this process to 92 grayscale natural images (Kriegeskorte et al., 2008). Reconstructed images were then used as input to a supervised AlexNet (Krizhevsky et al., 2012) (Figure 1) pretrained on the ImageNet object recognition dataset (Deng et al., 2009). For Figure 6, we used 10° d.v.a. reconstructions of the camel versus wrench familiar training stimuli for all model representations.

### Stimulus Presentation

Behavioral experiments were conducted in custom-built boxes connected to the animals’ home cages, allowing free movement between the cage and testing area. Infrared sensors recorded central and lateral nose pokes used for trial initiation and choice responses. Stimuli were presented on LCD monitors positioned behind a clear acrylic panel. Experimental events were controlled by a Mac Mini interfaced with two Arduino boards running MWorks software (https://mworks.github.io/). Images were displayed on a Feelworld T7 7” monitor with 1920 x 1200 resolution and 30Hz refresh rate at 3 inches of viewing distance.

### Behavioral Tasks

#### Camel vs. wrench and camel vs. rhino tasks

Before testing, animals were acclimated to the behavioral setup and trained to initiate trials. All three tree shrews (Se, Do, Ry) were first trained on a *camel versus wrench* discrimination task, which served as the baseline for subsequent experiments. Training began with a single target and distractor, with the camel always designated as the target. As performance stabilized, additional target and distractor images were gradually introduced until the full stimulus set was reached.

Each trial began when the animal initiated a central nose poke (100–450 ms hold duration). A reference camel image then appeared at the center of the screen, followed by two choice images presented subtending 10° d.v.a. Selecting the camel image triggered a juice reward (80% apple juice, 20% water). Incorrect responses resulted in a timeout period before the next trial, with the duration adjusted across animals (0–2700 ms). The median number of trials completed per session across all shrews was 864 (Supplementary Figure 2).

Once performance plateaued, animals entered a testing phase designed to assess generalization across untrained transformations. Each test session included both *novel rewarded* and *novel unrewarded* stimuli. Novel rewarded trials consisted of previously unseen transformations of the trained object identities and were rewarded for correct responses, providing a measure of generalization under reinforcement. Novel unrewarded trials consisted of additional untrained transformations that were never rewarded, preventing within-session learning. To prevent confusion caused by unrewarded trials, we periodically included reinforcement sessions containing two pairs of novel rewarded stimuli. This helped stabilize performance and maintain motivation across testing. For performance analysis, only test sessions containing both novel rewarded and novel unrewarded stimuli were used.

After completion of testing on the camel versus wrench familiar task, two animals (Se and Do) were trained and tested on a “unique” *camel versus wrench* variant in which the object categories were unchanged but the specific object exemplars differed from those used in the familiar condition, for example single- versus double-humped camels and box versus crescent wrenches. In this task variant, the original training set of “familiar” camels and wrenches were used as the training set and the “unique” camels and wrenches were used as the testing (*novel rewarded* and *novel unrewarded*) stimuli. This paradigm helped to keep tree shrews motivated with high performance on the training stimuli while assessing generalization across object exemplars within a category, rather than across transformations of the same exemplar.

In a separate task variant designed to increase shape similarity between target and distractor, two animals (Ry and Do) were trained and tested on a *camel versus rhino* discrimination. All details were consistent with the *camel versus wrench familiar* task besides the new distractor identity. One animal (Ry) completed both training and testing, whereas the second (Do) only completed training due to age-related health complications that affected behavioral performance.

#### Camel vs. wrench with backgrounds

To assess recognition under naturalistic conditions, the same animals were tested on a *camel versus wrench* task with objects embedded in natural scene backgrounds. Task structure and trial timing were identical to the earlier version, but each object transformation (three per category) was paired with six scene backgrounds shared targets and distractors. Each session presented eight image pairs (two transformations by four backgrounds), and transformation–background combinations were rotated across sessions until all pairings were shown. All trials were rewarded if tree shrews made correct nose pokes towards the image containing a camel (target).

### Behavior Analysis

#### Accuracy

Analyses of *camel versus wrench familiar, camel versus wrench unique,* and *camel versus rhino* tasks included only sessions with both *novel rewarded* and *novel unrewarded* stimuli. For the *camel versus wrench with backgrounds* experiment, sessions were included if more than four target images were presented. Across all tasks, sessions were excluded if average catch-trial performance was below 80% or if the session contained fewer than 100 total trials. For analyses focusing on specific target or target–distractor pairs, stimuli were excluded when fewer than ten trials were available.

#### Correlation Analyses

We examined both within-animal and between-animal correlations in behavioral performance to assess consistency and shared structure across tasks. All correlation analyses excluded target–distractor pairs with fewer than ten trials. Within-animal reliability was evaluated using correlations across 1,000 random splits of the trials for remaining target–distractor pair labels to measure internal consistency across splits. Null distributions were generated by shuffling target–distractor pairs, and correlations were considered significant if the observed mean exceeded the 97.5th percentile of the null distribution. Between-animal similarity was assessed by correlating 1,000 bootstrapped samples of target–distractor pairs between animals. In tasks with more than two tree shrews, correlations were between a single shrew’s target–distractor performance pattern and the average of the remaining two. Null distributions were generated by shuffling target–distractor pair labels, and significance was determined using the same 97.5th percentile threshold.

#### Background Contrast Analysis

To quantify the impact of low-level background statistics, like contrast, on behavior, we analyzed the root mean square (RMS) value for each of the six backgrounds in the absence of objects. We computed the average performance across all three shrews for all trials when targets and distractors were displayed on the same background. Since there were three unique target and distractor objects, this resulted in nine target-distractor pairs for each background. We then computed a line of best fit (Figure 5C) and Pearson’s correlation between the background RMS and shrew performance.

### Modeling of Tree Shrew Behavior

#### Stimulus Model Representations

All models shown in Figure 6 were computed using the training set from the camel versus wrench familiar condition. Stimuli were reconstructed using ISETTreeShrew at 10° d.v.a. For each model, we computed Euclidean distances between all target-distractor pair model representations and correlated these distances with the average behavioral performance across all shrews for the corresponding stimuli during test sessions.

#### Pixel-based representations

We first quantified similarity using pixel-level representations of the original stimuli (“*object*”). Because tree shrews were freely moving during the task, they may have partially compensated for changes in object position or size through head or body movements. To account for this possibility, we also generated position- and size-normalized versions of each stimulus.

Position normalization was performed by translating each object so that its center of mass aligned with that of the reference camel (“*object centered*”). Size normalization was performed by scaling each object so that its largest dimension (height or width) matched that of the reference camel (“*object scaled*”). If scaling caused the object to exceed image boundaries, the object was minimally translated to remain within bounds while preserving position information as much as possible. A combined normalization was also applied in which objects were first centered then scaled using the above procedures (“*object centered and scaled*”).

We also generated rotation-normalized images to account for the possibility that shrews compensate for in-plane image rotation through head tilt. However, since rotating the objects in the image plane does not change pixel values and rarely impacts target–distractor overlap, it minimally impacts distances between targets and distractors and is almost perfectly correlated with the original *object* representations (Pearson’s correlation of original and rotation-normalized image paired distances, r(82) = 0.96, r > 97.5 percentile of null distribution from 1000 random permutations). Rotation-normalized images were therefore excluded from further analyses. Euclidean pixel distances between all target-distractor pairs were computed for each representation and compared with behavioral performance.

#### Feature-based representations

To assess the contribution of progressively more complex visual features, we generated several feature-based stimulus representations intended to approximate different stages of hierarchical visual processing.

For the *Gabor* model, we applied Matlab’s imgaborfilt using eight spatial frequencies (5- 75 pixels/cycle) and twelve orientations (0-165°), yielding 96 filters. For each stimulus, filter responses were averaged across pixels to yield a single response value per filter. Each filter was z-scored across stimuli (separately for targets and distractors) to eliminate magnitude differences across filter types. Euclidean distances between these filter response vectors were then computed for all target–distractor pairs.

Texture representations were generated using Plenoptic’s (Balzani et al., 2023; Duong et al., 2023) implementation of a texture model (Portilla and Simoncelli, 2000). Two texture variants were created. In the first texture model (“*texture*”), stimuli were cropped using a square bounding box so that the object filled most of the image and then resized back to the original dimensions prior to metamer synthesis. In the second texture model (“*texture-in-place*”), we sought to create a texture while preserving the object’s size and position information. Objects were cropped using rectangular bounding boxes that closely matched object boundaries while remaining large enough for processing (minimum 64 pixels per dimension). The resulting texture was then placed back into its original spatial position on a uniform background set to the mean texture luminance. For both texture models, we used the MetamerCTF procedure with two scales, a maximum of 1500 iterations, and an L2 norm loss function.

*Saliency* representations were computed using SALICON (Jiang et al., 2015), which is trained on human eye tracking annotations from the Microsoft Common Objects in Context (MS COCO; Lin et al., 2015) image database. Saliency maps were inverted to match the luminance polarity of other model representations.

*Binary* representations were generated by binarizing the image based on the background value, then removing any small holes in the object with an area threshold of 3 pixels (Walt et al., 2014). Following this, we slightly expanded the object with a kernel of [3,3] to maintain a smooth edge using OpenCV’s image processing tools (Bradski, 2000). The *skeleton* representations were then generated from these binary images by extracting the medial-axis skeleton using skeletonize from scikit-image (Lee et al., 1994), and applying a Gaussian blur with a kernel width of [5,5] pixels to smooth the skeleton. Skeleton images were also inverted to match luminance polarity across models. *Binary object, scaled* and *skeleton scaled* representations were made by applying the same transformations of the *object scaled* images.

All stimulus transformation methodologies excluding SALICON can be found on GitHub (github.com/emeyer121/StimulusModeling).

#### Feature abstraction via shallow convolutional models

For the texture, saliency, binary, and skeleton models, we implemented an additional abstraction step to reduce sensitivity to precise spatial position and emphasize feature content. For each representation type, we trained a shallow convolutional neural network consisting of a single convolutional layer followed by ReLU activation, pooling, and a linear projection. Networks were trained to classify targets versus distractors using binary cross-entropy loss with logits (BCEWithLogitsLoss). Training used 88 target and 93 distractor images that excluded the 12 experimental targets and 7 distractors, totaling 100 targets and 100 distractors. Data were split into five folds, and models were trained until validation accuracy plateaued (50-350 epochs) with batch size of 32 and a learning rate of 0.005. The weights across folds were averaged to form an ensemble model for each representation type. Experimental stimuli were then passed through the ensemble model to extract activations. Test accuracies on held out images varied across representations, ranging from 0.53-1. Euclidean distances between activations for each target-distractor pair were computed and compared with behavioral performance.

#### Between-Model Correlations

To assess overlap between different model representations, we computed a correlation matrix by calculating Pearson correlations between target-distractor distance values for all pairs of models (Supplementary Figure 6). Using this correlation matrix, we computed hierarchical clustering using the average linkage algorithm which resulted in two clusters of four models each (Cluster 1, green; Cluster 2, blue) with the remaining models unclustered (gray). We then computed correlation of each model’s target-distractor distances with behavior, organized based on the hierarchical clustering (Figure 6A). Error was computed using 1000 bootstrapped samples and correlations were significant if 95% confidence intervals exceeded zero.

#### Convolutional neural network models

To compare tree shrew behavior with standard hierarchical visual representations commonly used in studies of object recognition, we analyzed three deep neural network models: a supervised AlexNet (Krizhevsky et al., 2012), a self-supervised contrastive AlexNet trained on lower-resolution ImageNet images (Nayebi et al., 2023), and a supervised VGG16 (Simonyan and Zisserman, 2015) all pre-trained on ImageNet (Deng et al., 2009). For both AlexNet models, we extracted activations from the max pooling or ReLU layers preceding convolutional layers to make up our set of “Conv” layers. Since VGG16 has additional convolutional layers, we extracted activations from each max pooling layer in order to match the number of “Conv” layers from other models. For all three CNNs, linear layer activations were extracted to make up the set of fully connected (fc) layers (3 for supervised AlexNet and VGG16, 2 for contrastive AlexNet). Euclidean distances between activations at each layer for all target-distractor pairs were then computed and compared with behavioral performance. Error was computed using 1000 bootstrapped samples and correlations were marked as significant if 95% confidence intervals exceeded zero (Figure 6B).

Knowing that object size is a confounding factor in behavioral performance, we also computed correlations of neural networks with behavior after regressing out target-distractor size differences from the correlation (Figure 6C). For CNN correlations, we used the final fc layer from each model. Error bars were computed using 1000 bootstrapped samples and correlations were significant if 95% confidence intervals exceeded zero.

## Acknowledgements

We thank N Rust, B Jannuzi, and T Meyer for work on initial versions of ISETTreeShrew and tree shrew behavioral piloting, as well as A Roorda for providing data on tree shrew optics in tabular form.

## Extended Data

**Supplementary Figure 1:**
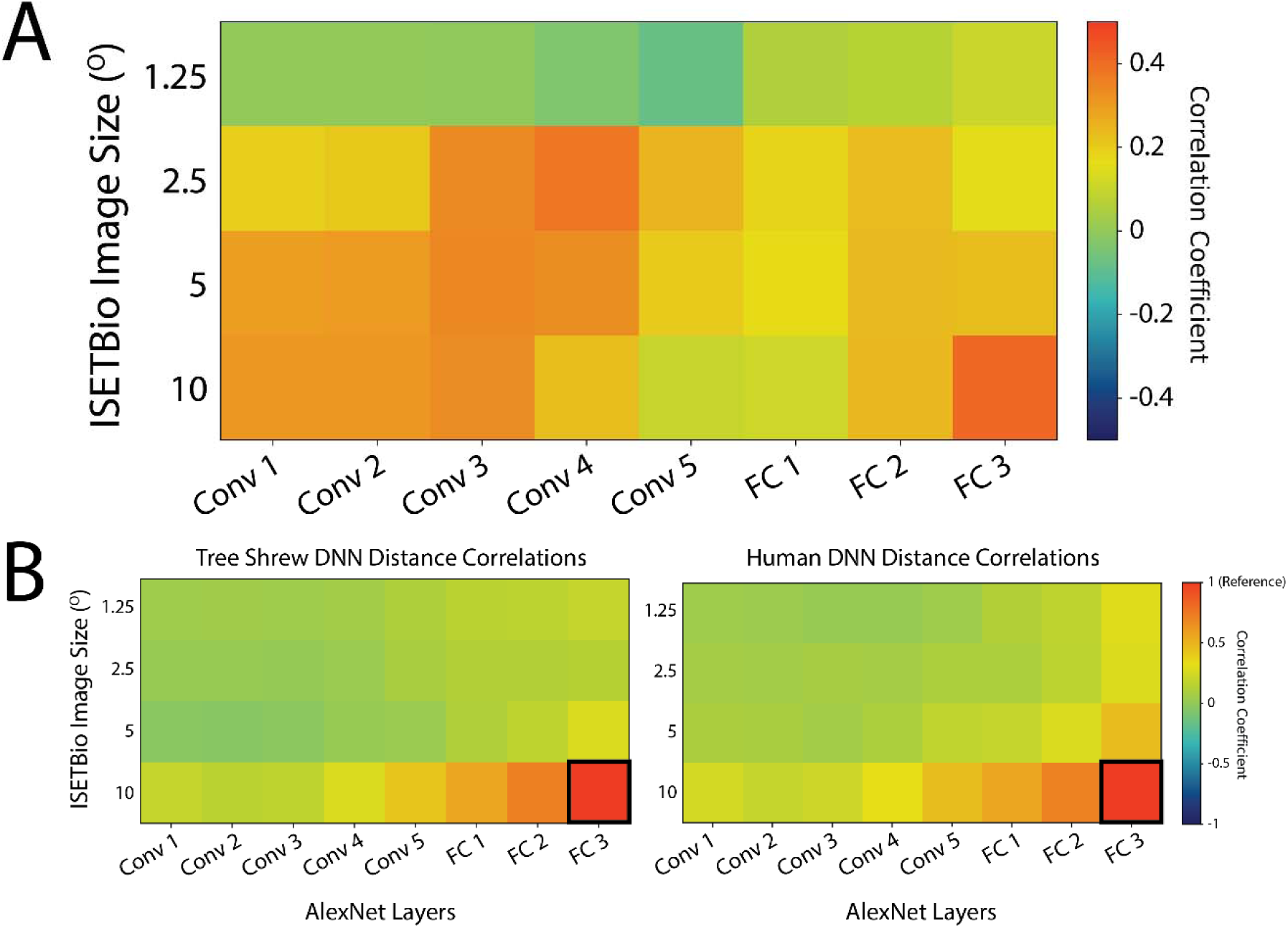
DNN RDM correlations of tree shrew and humans across image sizes. (A) Pearson correlations of activation patterns in response to images filtered by human and tree shrew optics. Values at each size and layer correspond to the correlations of human and tree shrew RDMs of Euclidean distances between all image pair AlexNet activations at a specific layer and simulated image size. Images are grayscaled versions of 92 objects from Kriegeskorte et al., 2008. (B) Using same RDMs of activation distances, each square represents the within-species correlation with the “reference” size and layer of a 10° image in AlexNet layer FC3 (left, tree shrew; right, human).

**Supplementary Figure 2:**
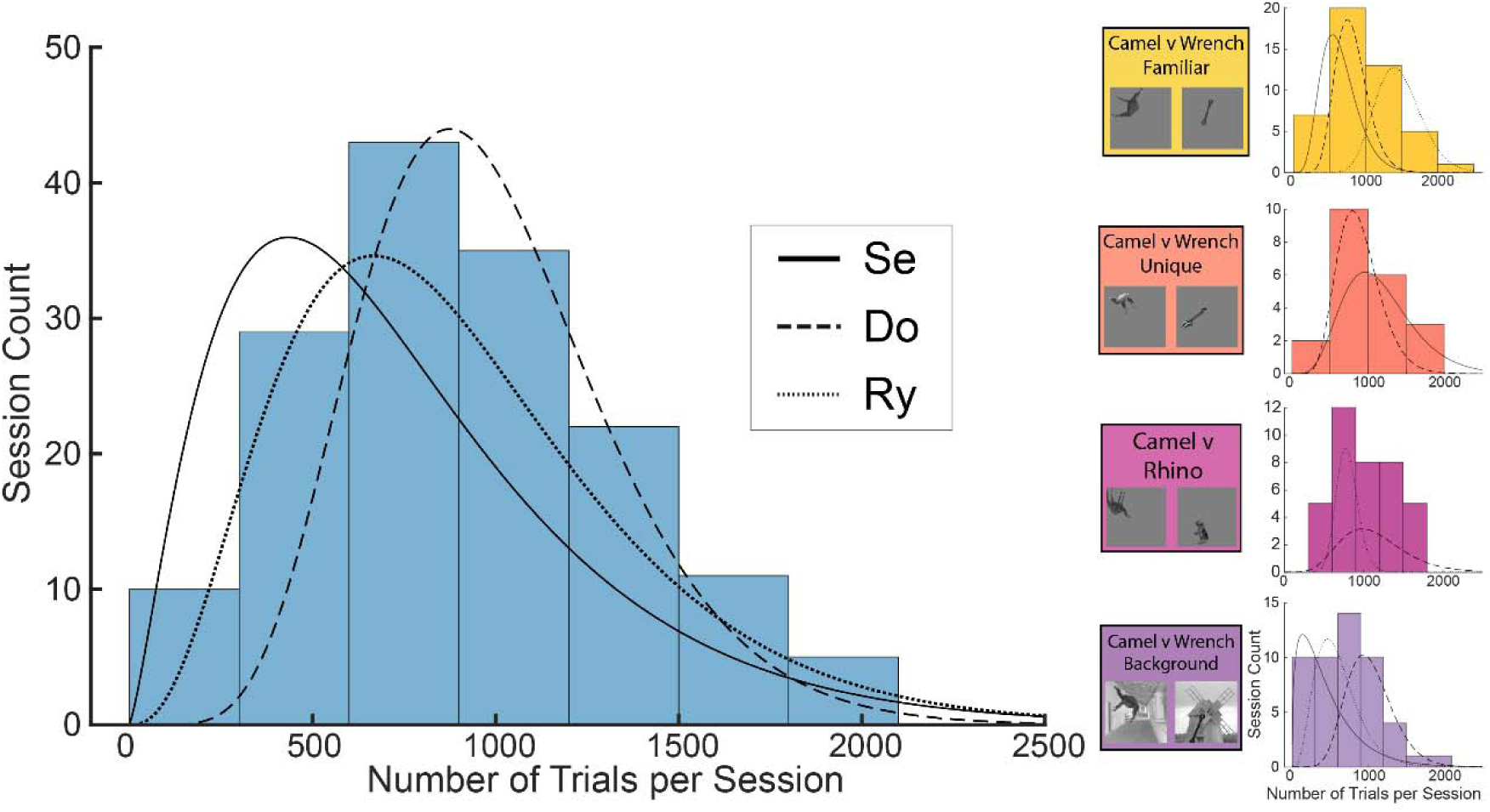
Distribution of trial counts per session from all shrews and tasks. Histogram of number of trials performed in each session across all shrews and all experiment versions (left). Median value is 864 trials. Curves represent negative binomial fits for each individual shrew trial distributions. Figures on the right depict number of trials performed per session in individual experiment types.

**Supplementary Figure 3:**
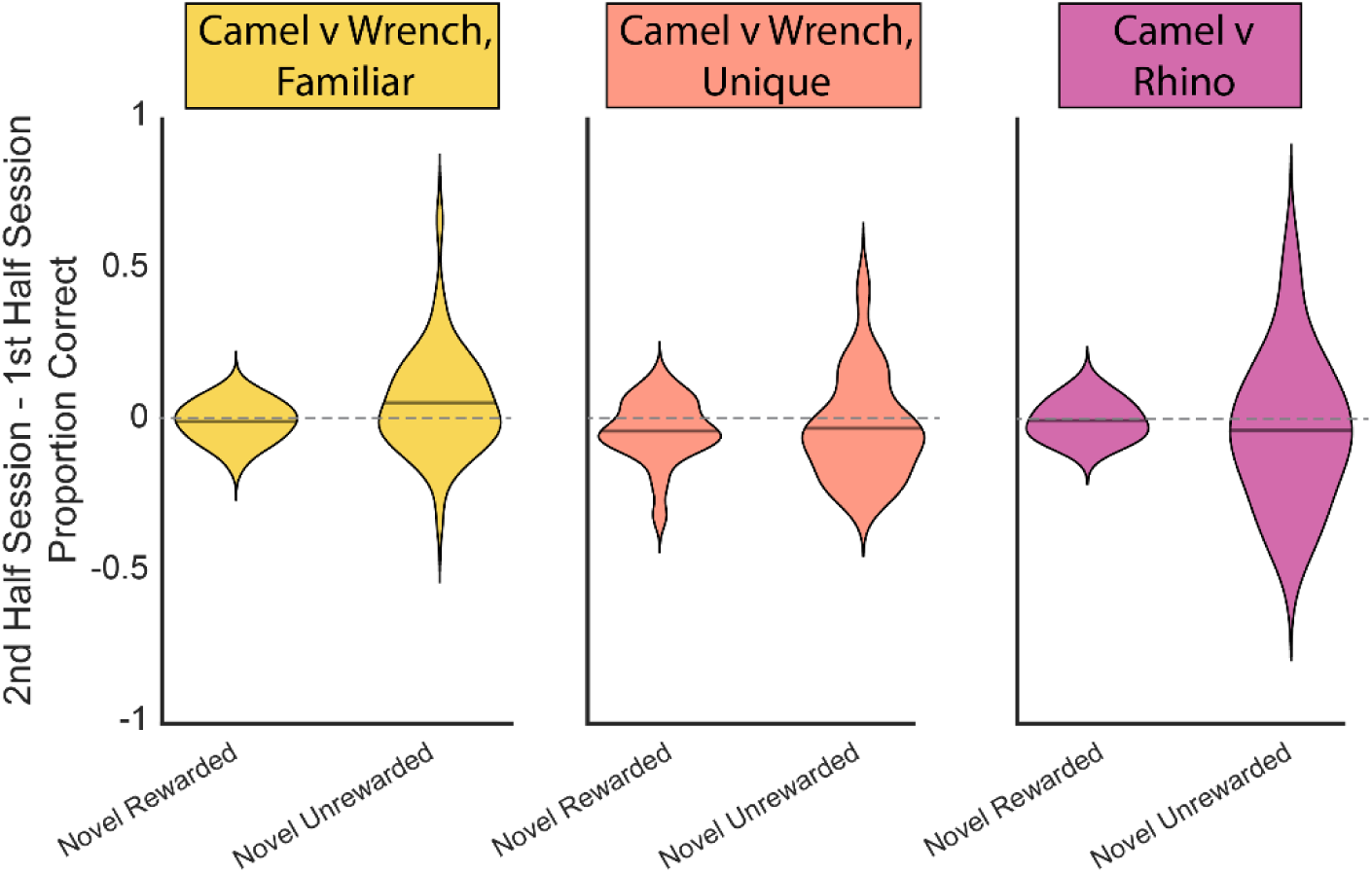
Violin plots depicting average performance difference on novel rewarded and unrewarded stimuli between first and second half of sessions. Performance values are collapsed across all shrews in each experiment type. Solid horizontal line in violin plot represents mean value.

**Supplementary Figure 4:**
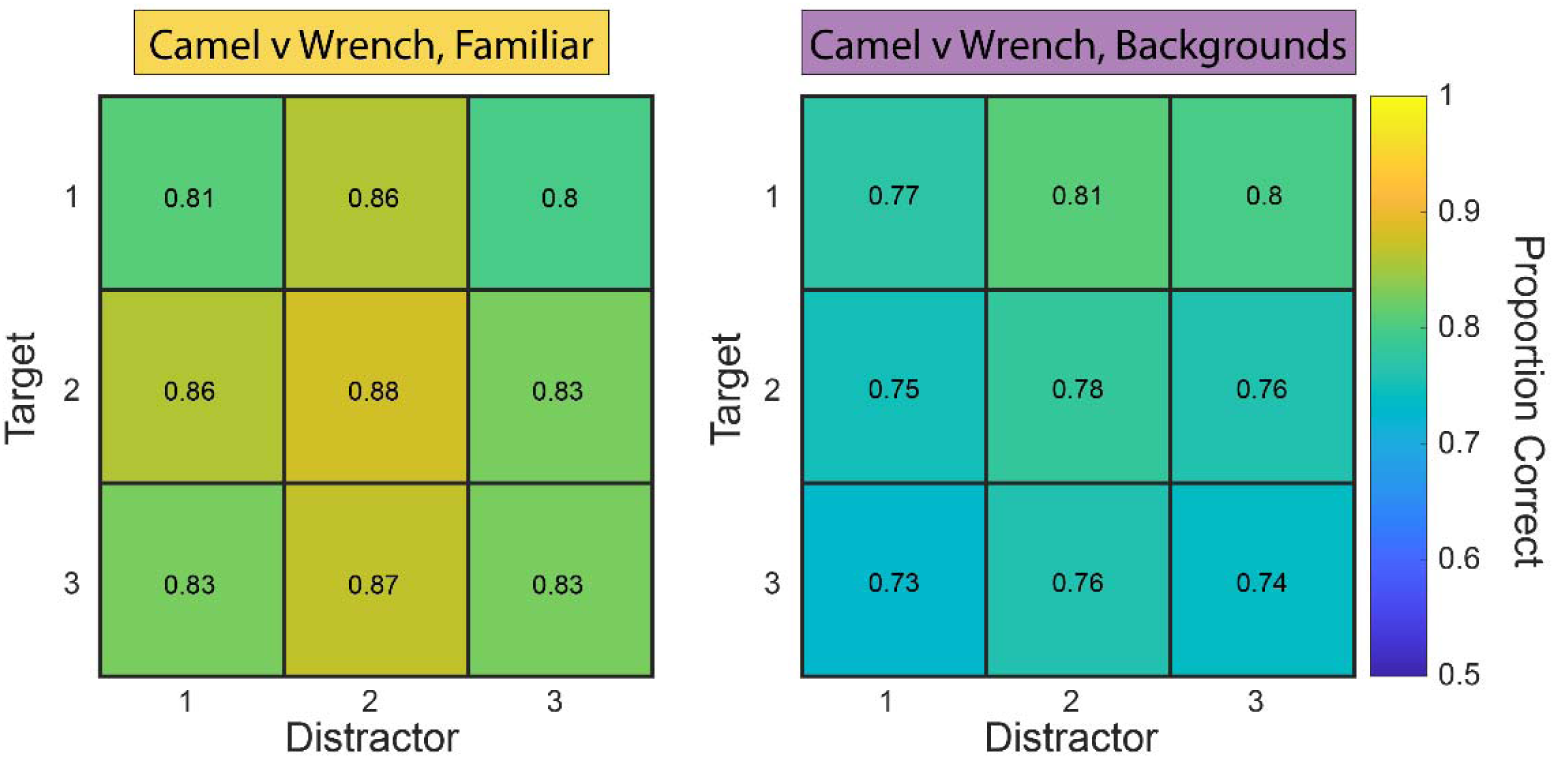
Comparison of target-distractor performance with and without backgrounds. Average performance collapsed across shrews for each target-distractor pair presented in camel versus wrench with backgrounds task and performance on corresponding stimuli presented on mean gray backgrounds. Pearson correlation between performance values is r(7) = -0.09, p = 0.82.

**Supplementary Figure 5:**
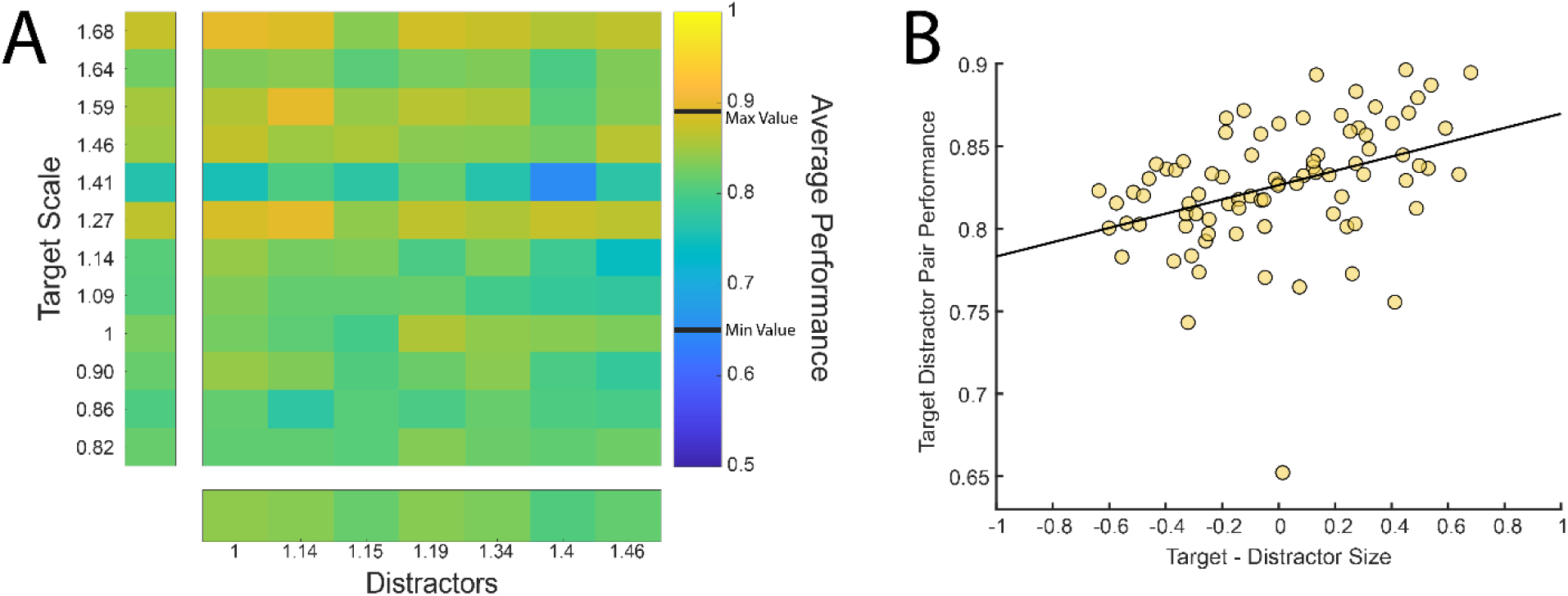
Relationship of target and distractor size with discrimination performance. (A) Average performance for all shrews on training set of images in camel versus wrench familiar task. Target and distractor objects are ordered based on scale factor. (B) Correlation between average target-distractor pair performance and difference in corresponding scale factors. Pearson correlation between performance and size is r(82) = 0.39, p = 0.00026.

**Supplementary Figure 6:**
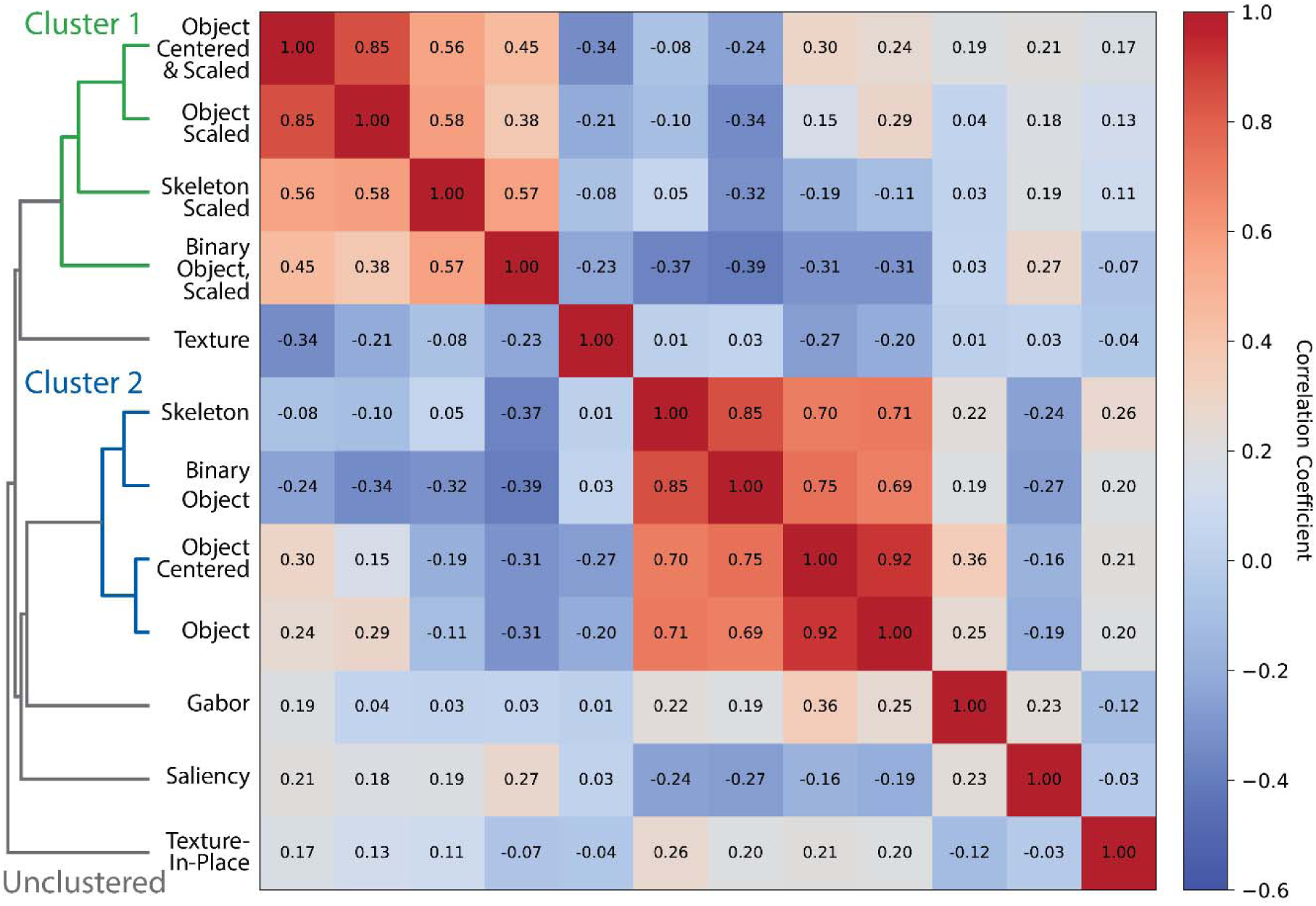
Hierarchical clustering of model relationships. Correlation matrix based on pattern of camel versus wrench familiar target-distractor distances for each model type. Models are organized based on hierarchical clustering using the average linkage algorithm.

**Supplementary Figure 7:**
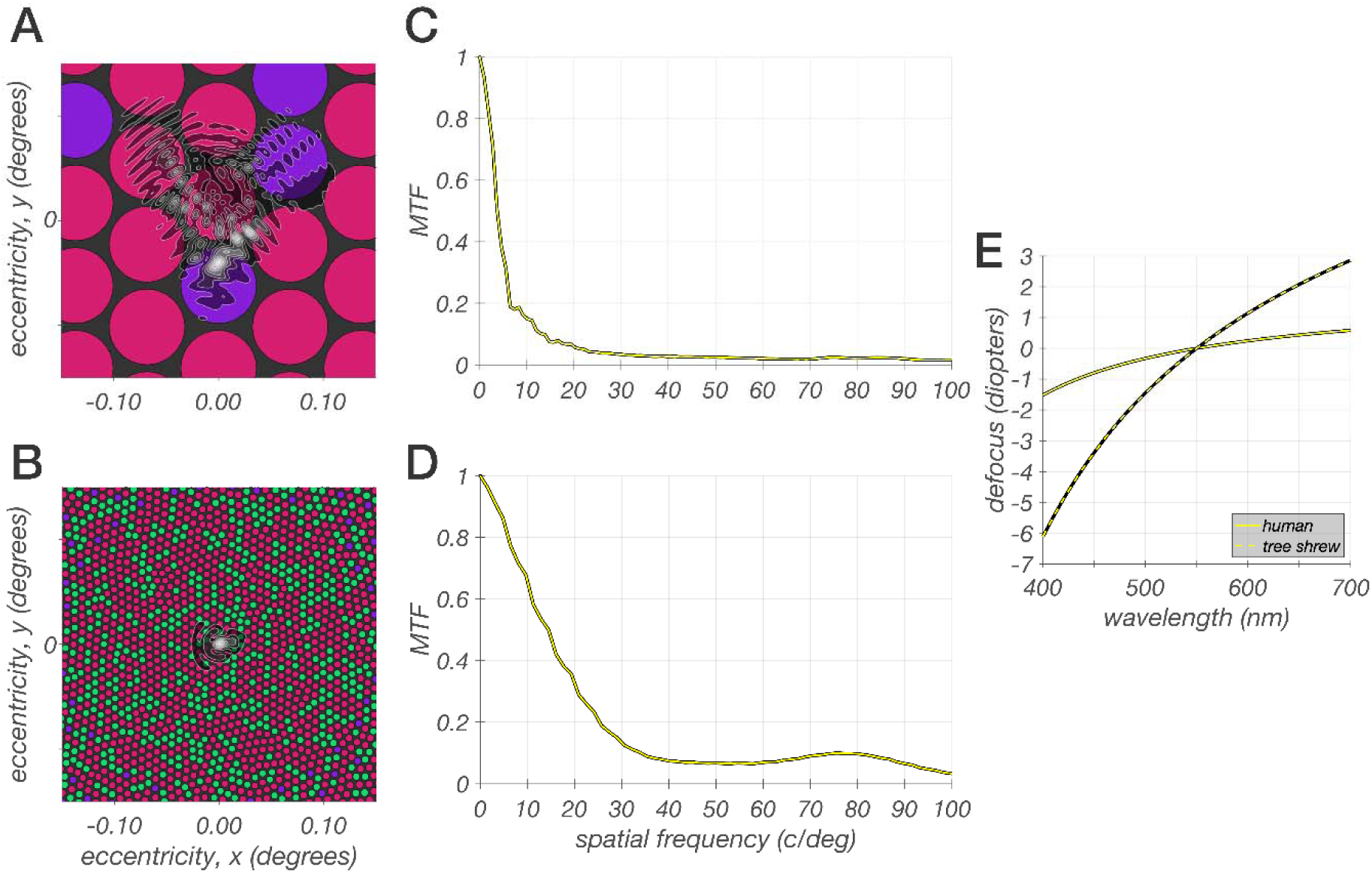
ISETBio model of the tree shrew initial visual encoding. (A) ISETBio model of a treeshrew cone mosaic model spanning a region of 0.3 x 0.3 degrees. L and S-cones are depicted by the red and purple disks. The superimposed contour plot depicts the ISETBio point spread function (PSF) of a tree shrew at 550 nm and a 4.0 mm pupil for a typical subject from the study of Sajdak et al., 2019. (B) ISETBio model of a human cone mosaic model spanning a region of 0.3 x 0.3 degrees. L, M, and S-cones are depicted by the red, green and purple disks. In the human cone mosaic, there are no S-cones in the central 0.3 degrees. The superimposed contour plot depicts a human PSF at 550 nm and a 4.0 mm pupil for a typical subject from the study of Polans et al., 2015. Note change in scale relative to A. (C) Radially averaged modulation transfer function (MTF) of the treeshrew point spread function depicted in A. At around 7 c/deg, the optics alone reduce retinal contrast by 80% and virtually eliminate it for spatial frequencies above 30 c/degs. Further significant contrast reduction at the level of the cone excitations is introduced due to blurring by the large light gathering apertures of three shrew cones (effect not shown). (D) Radially averaged MTF of the human point spread function depicted in B. At around 25 c/deg, the human optics reduce retinal contrast by 80% and virtually eliminate it for spatial frequencies above 100 c/degs. Further contrast reduction at the level of the cone excitations is introduced due to blurring by the cone apertures (effect not shown), but this effect is smaller than in tree shrew. (E) The additional defocus of retinal contrast as a function of wavelength, which introduces longitudinal chromatic aberration, is depicted by the solid curve for the human eye, and by the dotted curve for the tree shrew eye. Note that over the visible spectrum there is about 2.5 diopters of defocus in the human eye, and around 9 diopters of defocus in the tree shrew eye. Note however, that power of the optics of the two eyes is quite different, so that the effect of wavelength-dependent focus is calculated in an eye specific manner.

