## Supplemental Figures for "Transformation-tolerant object recognition in tree shrews despite lacking a fovea"

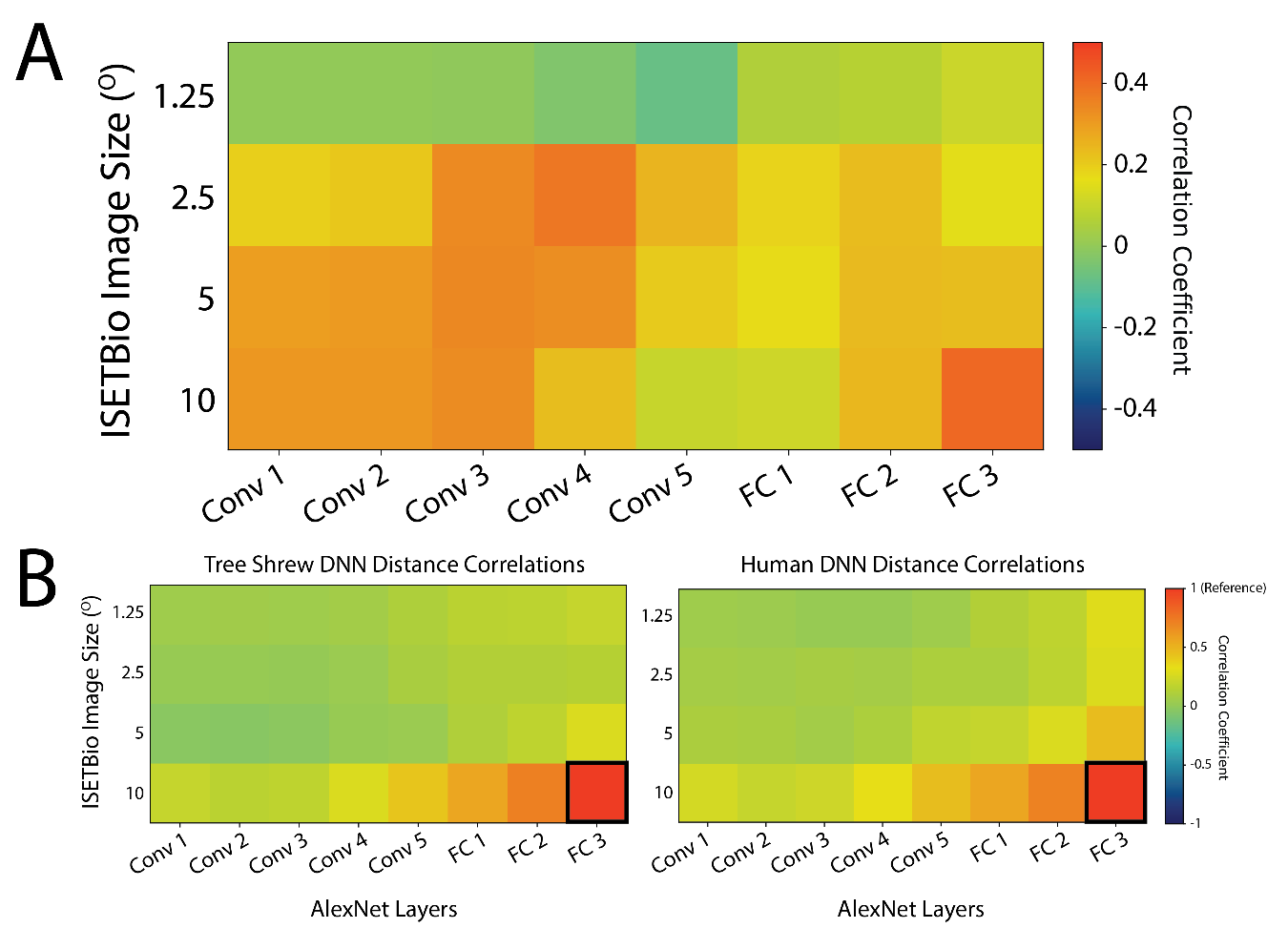

**Supplementary Figure 1: DNN RDM correlations of tree shrew and humans across image sizes.** (A) Pearson correlations of activation patterns in response to images filtered by human and tree shrew optics. Values at each size and layer correspond to the correlations of human and tree shrew RDMs of Euclidean distances between all image pair AlexNet activations at a specific layer and simulated image size. Images are grayscaled versions of 92 objects from Kriegeskorte et al., 2008. (B) Using same RDMs of activation distances, each square represents the within-species correlation with the “reference” size and layer of a 10° image in AlexNet layer FC3 (left, tree shrew; right, human).

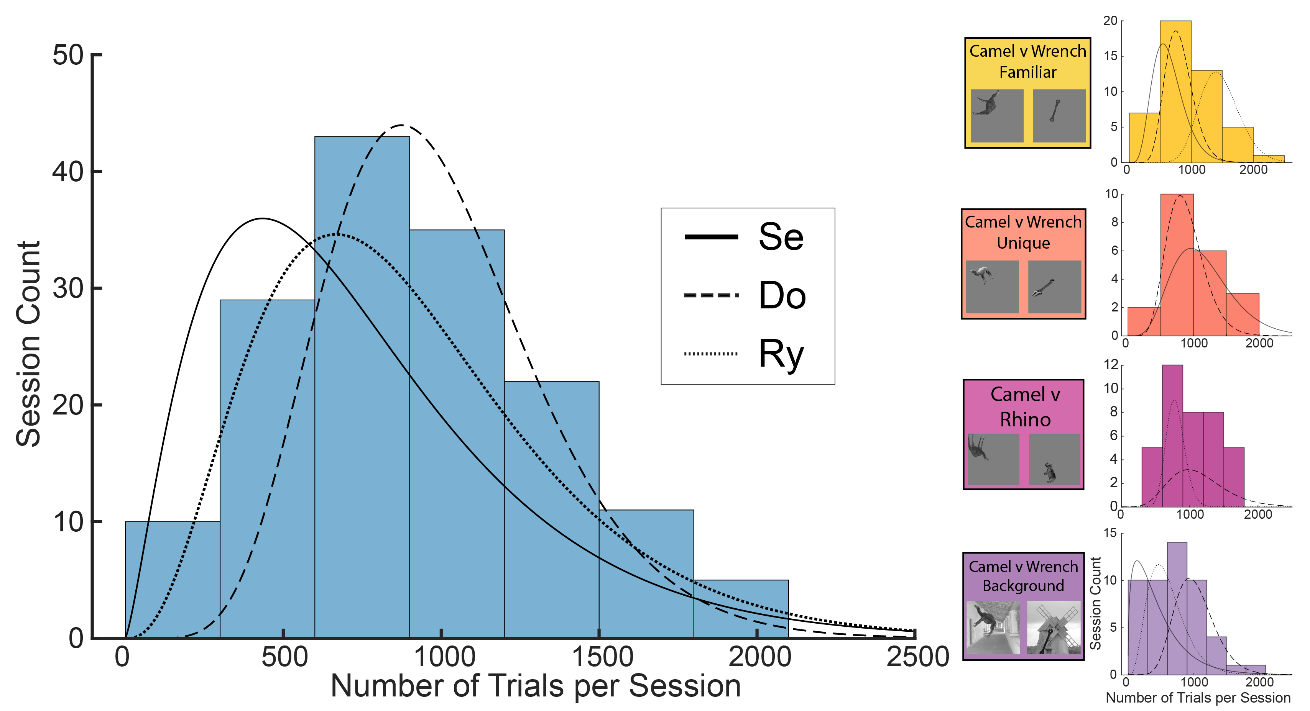

**Supplementary Figure 2: Distribution of trial counts per session from all shrews and tasks.** Histogram of number of trials performed in each session across all shrews and all experiment versions (left). Median value is 864 trials. Curves represent negative binomial fits for each individual shrew trial distributions. Figures on the right depict number of trials performed per session in individual experiment types.

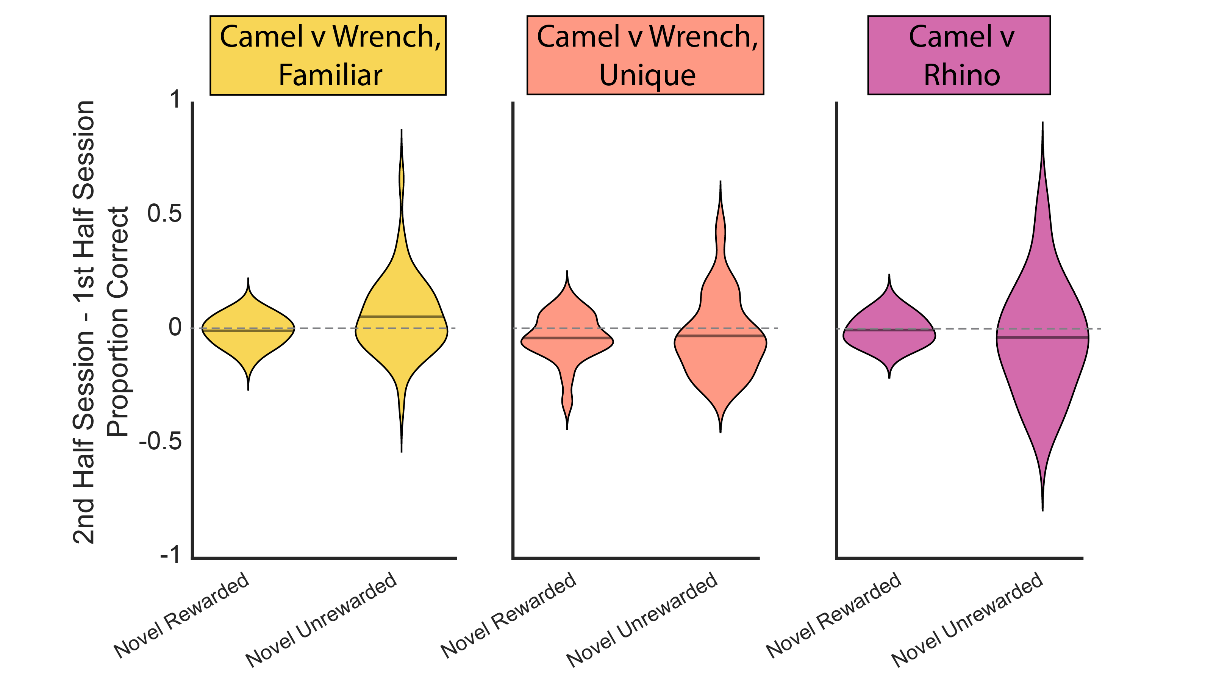

**Supplementary Figure 3:** Violin plots depicting average performance difference on novel rewarded and unrewarded stimuli between first and second half of sessions. Performance values are collapsed across all shrews in each experiment type. Solid horizontal line in violin plot represents mean value.

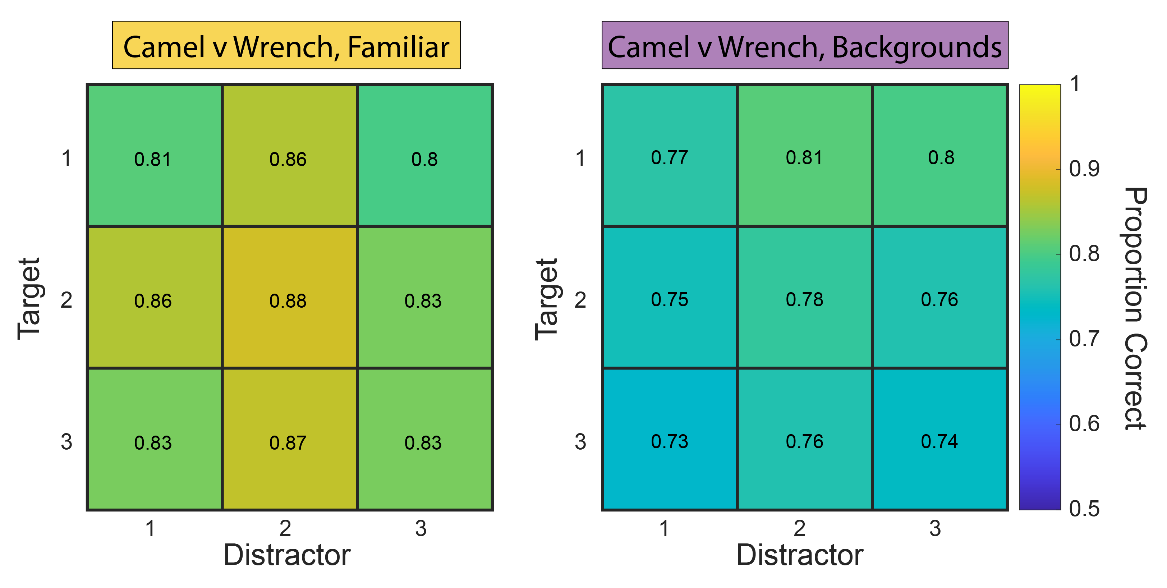

**Supplementary Figure 4: Comparison of target-distractor performance with and without backgrounds.** Average performance collapsed across shrews for each target-distractor pair presented in camel versus wrench with backgrounds task and performance on corresponding stimuli presented on mean gray backgrounds. Pearson correlation between performance values is r(7) = -0.09, p = 0.82.

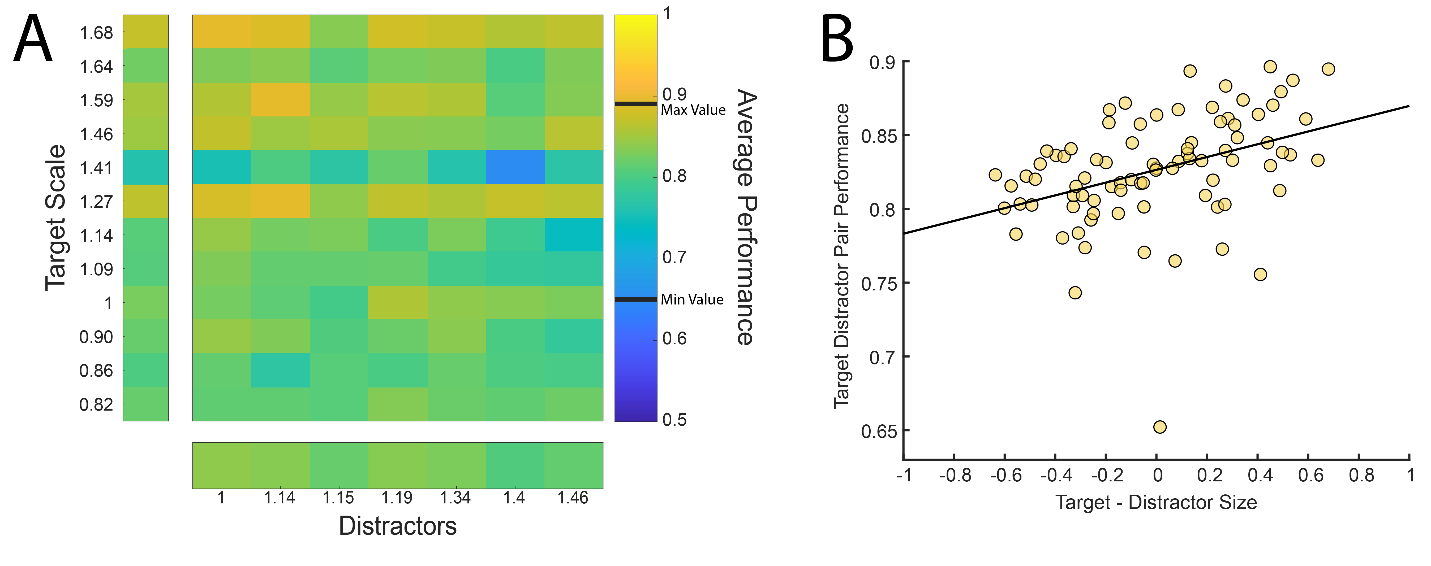

**Supplementary Figure 5: Relationship of target and distractor size with discrimination performance.** (A) Average performance for all shrews on training set of images in camel versus wrench familiar task. Target and distractor objects are ordered based on scale factor. (B) Correlation between average target-distractor pair performance and difference in corresponding scale factors. Pearson correlation between performance and size is r(82) = 0.39, p = 0.00026.

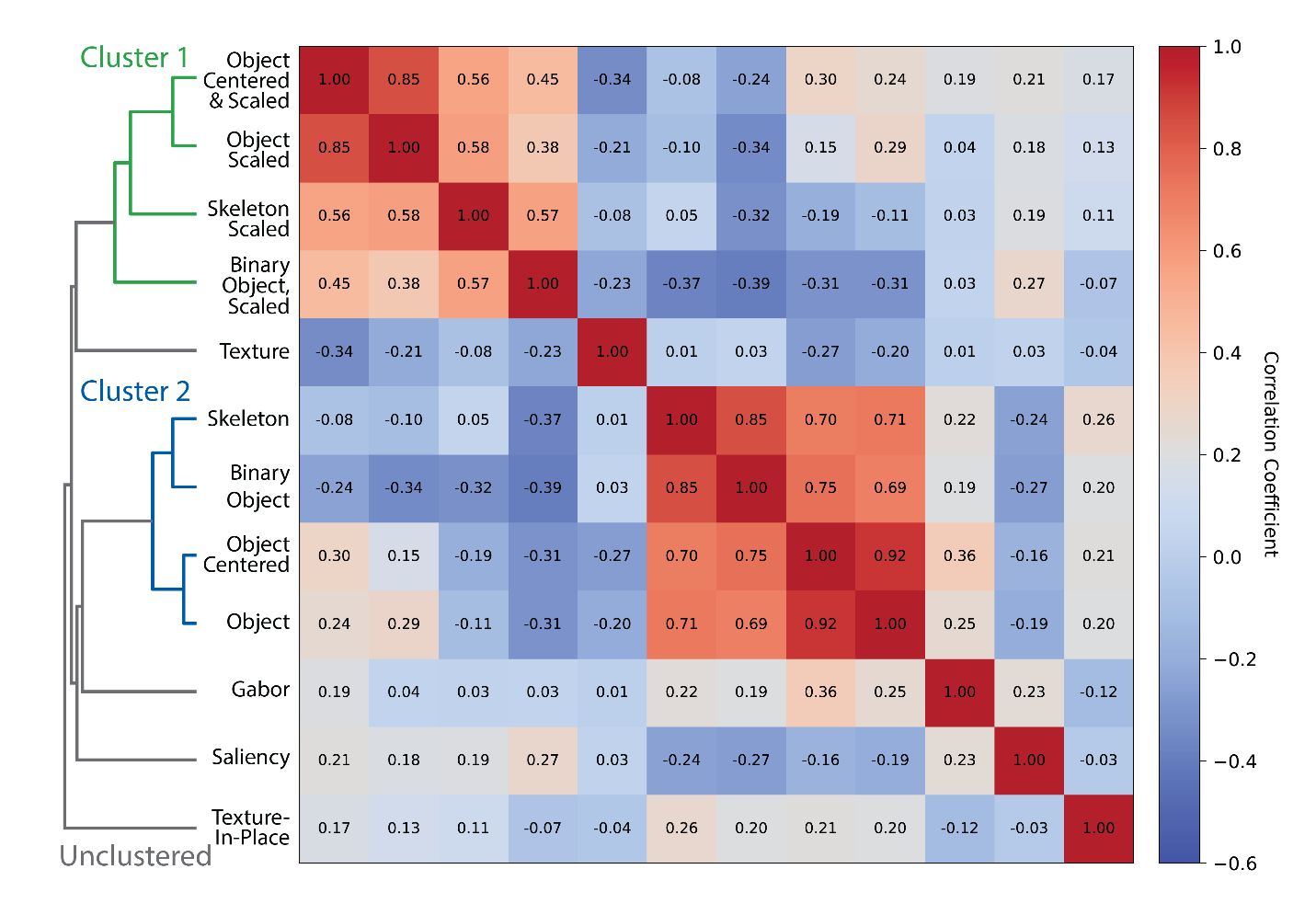

**Supplementary Figure 6: Hierarchical clustering of model relationships.** Correlation matrix based on pattern of camel versus wrench familiar target-distractor distances for each model type. Models are organized based on hierarchical clustering using the average linkage algorithm.

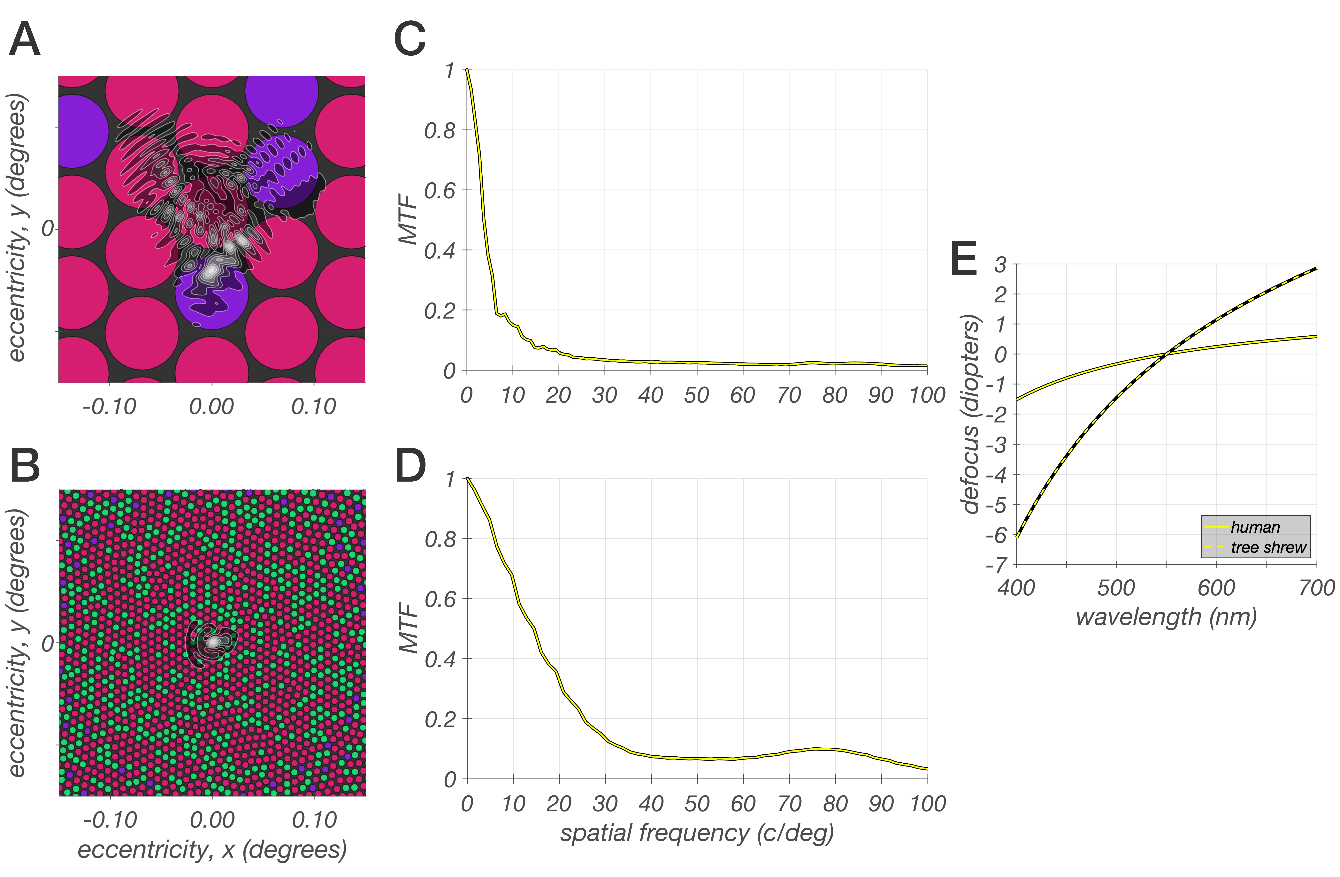

**Supplementary Figure 7: ISETBio model of the tree shrew initial visual encoding.** (A) ISETBio model of a treeshrew cone mosaic model spanning a region of 0.3 x 0.3 degrees. L and S-cones are depicted by the red and purple disks. The superimposed contour plot depicts the ISETBio point spread function (PSF) of a tree shrew at 550 nm and a 4.0 mm pupil for a typical subject from the study of Sajdak et al., 2019. (B) ISETBio model of a human cone mosaic model spanning a region of 0.3 x 0.3 degrees. L, M, and S-cones are depicted by the red, green and purple disks. In the human cone mosaic, there are no S-cones in the central 0.3 degrees. The superimposed contour plot depicts a human PSF at 550 nm and a 4.0 mm pupil for a typical subject from the study of Polans et al., 2015. Note change in scale relative to A. (C) Radially averaged modulation transfer function (MTF) of the treeshrew point spread function depicted in A.  At around 7 c/deg, the optics alone reduce retinal contrast by 80% and virtually eliminate it for spatial frequencies above 30 c/degs. Further significant contrast reduction at the level of the cone excitations is introduced due to blurring by the large light gathering apertures of three shrew cones (effect not shown). (D) Radially averaged MTF of the human point spread function depicted in B. At around 25 c/deg, the human optics reduce retinal contrast by 80% and virtually eliminate it for spatial frequencies above 100 c/degs. Further contrast reduction at the level of the cone excitations is introduced due to blurring by the cone apertures (effect not shown), but this effect is smaller than in tree shrew. (E) The additional defocus of retinal contrast as a function of wavelength, which introduces longitudinal chromatic aberration, is depicted by the solid curve for the human eye, and by the dotted curve for the tree shrew eye. Note that over the visible spectrum there is about 2.5 diopters of defocus in the human eye, and around 9 diopters of defocus in the tree shrew eye. Note however, that power of the optics of the two eyes is quite different, so that the effect of wavelength-dependent focus is calculated in an eye specific manner.

**Camel versus wrench familiar**

| **Shrew Name** | **# Test Sessions Included** | **Avg Trials Per Sesion (+/- SD)** | **Exp Perf (+/- SD)** | **Catch Perf (+/- SD)** | **Within-Shrew Corr (95% CI)** |
| --- | --- | --- | --- | --- | --- |
| Se | 15 | 643 (+/- 265) | 0.84 (+/- 0.038) | 0.92 (+/- 0.039) | 0.27 (0.16-0.38) |
| Do | 15 | 809 (+/- 209) | 0.83 (+/- 0.031) | 0.89 (+/- 0.037) | 0.43 (0.34-0.53) |
| Ry | 16 | 1463 (+/- 332) | 0.79 (+/- 0.045) | 0.87 (+/- 0.037) | 0.24 (0.17-0.31) |

**Camel versus wrench novel**

| **Shrew Name** | **# Test Sessions Included** | **Avg Trials Per Sesion (+/- SD)** | **Exp Perf (+/- SD)** | **Catch Perf (+/- SD)** | **Within-Shrew Corr (95% CI)** |
| --- | --- | --- | --- | --- | --- |
| Se | 11 | 1150 (+/- 413) | 0.82 (+/- 0.051) | 0.94 (+/- 0.033) | 0.54 (0.47-0.62) |
| Do | 10 | 884 (+/- 312) | 0.82 (+/- 0.025) | 0.87 (+/- 0.049) | 0.34 (0.24-0.44) |

**Camel versus rhino**

| **Shrew Name** | **# Test Sessions Included** | **Avg Trials Per Sesion (+/- SD)** | **Exp Perf (+/- SD)** | **Catch Perf (+/- SD)** | **Within-Shrew Corr (95% CI)** |
| --- | --- | --- | --- | --- | --- |
| Do | 24 (only training) | 1125 (+/- 366) | 0.67 (+/- 0.038) | 0.84 (+/- 0.026) | 0.68 (0.60-0.77) |
| Ry | 14 | 793 (+/- 136) | 0.79 (+/- 0.040) | 0.87 (+/- 0.028) | 0.32 (0.22-0.41) |

**Camel v wrench backgrounds**

| **Shrew Name** | **# Test Sessions Included** | **Avg Trials Per Sesion (+/- SD)** | **Exp Perf (+/- SD)** | **Catch Perf (+/- SD)** | **Within-Shrew Corr (95% CI)** |
| --- | --- | --- | --- | --- | --- |
| Se | 9 | 454 (+/- 534) | 0.73 (+/- 0.054) | 0.99 (+/- 0.019) | 0.60 (0.51-0.69) |
| Do | 17 | 1025 (+/- 314) | 0.72 (+/- 0.051) | 0.91 (+/- 0.038) | 0.56 (0.50-0.62) |
| Ry | 24 | 600 (+/- 274) | 0.66 (+/- 0.044) | 0.89 (+/- 0.045) | 0.47 (0.39-0.55) |

**Table 1: Summary statistics of task variations.** Specific details of each task variation including shrews, session and trial counts, performance and consistency metrics.
